# Engineered chitosan-derived nanocarrier for efficient siRNA delivery to peripheral and central neurons

**DOI:** 10.1101/2024.09.05.611434

**Authors:** Ana P. Spencer, Adriana Vilaça, Miguel Xavier, Rafael Santos, Ariel Ionescu, María Lázaro, Victoria Leiro, Eran Perlson, Sofia C. Guimarães, Ben M. Maoz, Ana P. Pêgo

## Abstract

Gene therapy using small interfering RNA (siRNA) holds promise for treating neurological disorders by silencing specific genes, such as the phosphatase and tensin homolog (*PTEN*) gene, which restricts axonal growth. Yet, delivering siRNA to neurons efficiently is challenging due to premature degradation and unspecific delivery. Chitosan-based delivery systems have shown great potential due to their well-established biocompatibility. However, their limited transfection efficiency and lack of neuronal tropism require further modification. Building on our previous successes with neuron-targeted DNA delivery using chitosan, a novel approach for siRNA delivery aimed at PTEN downregulation is proposed. This involves using thiolated trimethyl chitosan (TMCSH)-based siRNA nanoparticles functionalized with the neurotropic C-terminal fragment of the tetanus neurotoxin heavy chain (HC) for efficient delivery to both peripheral and central neurons. These polyplexes demonstrated suitable physicochemical properties, biocompatibility, and no adverse effects on neuronal electrophysiology. Diverse neuronal models, including 3D *ex vivo* cultures and microfluidics, confirmed polyplexes’ efficiency and neurospecificity. HC targeting significantly enhanced nanoparticle neuronal binding, and live cell imaging revealed five times faster retrograde transport along axons. Furthermore, siRNA delivery targeting PTEN promoted axonal outgrowth in embryonic cortical neurons. Thus, these polyplexes represent a promising platform for siRNA delivery, offering potential for clinical translation and therapeutic applications.

## 1. Introduction

Gene therapy with a specific focus on small interfering RNA (siRNA) is still in its infancy in the context of nervous system applications yet holds immense potential for revolutionizing the treatment landscape of several neurological disorders [1–3]. This innovative therapeutic strategy, which is actively being explored, involves the silencing of specific genes expression to address underlying genetic anomalies or dysregulations associated with neuronal diseases. One prominent target is the Phosphatase and Tensin Homolog (PTEN) encoding gene. Dysregulation of PTEN, which plays a crucial role in controlling cell growth, survival, and apoptosis, has been implicated in various neurodegenerative conditions, making it a prime candidate for neuronal therapeutic intervention [4–7].

The premise of siRNA-based gene therapy for neurons revolves around the precise silencing of PTEN expression to modulate key cellular processes perturbed in neurological diseases. This approach involves designing siRNA sequences that specifically target PTEN messenger RNA (mRNA) to be delivered to neuronal cells, enabling a selective downregulation of PTEN protein levels. The ultimate goal is to promote neuronal survival, enhance synaptic plasticity, and ameliorate the pathological processes associated with neurodegenerative disorders [4–7]. Nanoparticle (NP)-based approaches, particularly those constructed from biocompatible polymers like chitosan, have garnered significant attention [8, 9]. Despite the promise of this therapy, its success hinges on overcoming several challenges associated with NP development. These issues include lack of neuronal tropism, cytotoxicity, and low transfection and downregulation efficiencies.

Chitosan, a naturally occurring heteropolysaccharide derived from chitin through N-deacetylation, has been extensively explored across various biomedical realms, gene delivery being a prominent focus [10–12]. Recognized as a promising gene carrier, chitosan boasts significant advantages in regenerative medicine, particularly its low cytotoxicity and capacity to mediate transient gene expression. Notably, the U.S. Food and Drug Administration (FDA) has approved its use in certain biomedical applications, emphasizing its safety and efficacy [13, 14]. Nevertheless, the limited transfection efficiency of chitosan under physiological conditions has posed a hurdle to its widespread implementation [15]. Furthermore, such vectors do not have any particular tropism in the nervous system. Nevertheless, the functional groups present in the chitosan backbone allow for modifications, facilitating the incorporation of ligands that can selectively bind to receptors expressed on neuronal cells that can ensure precise delivery to the intended neuronal targets.

Previously, our team has developed a suitable chitosan-based NP for the neuron-targeted delivery of therapeutic plasmid DNA (pDNA) to peripheral neurons [16–19]. In this pursuit, the material used was a modified chitosan derivative – partially thiolated trimethyl chitosan (TMCSH), actively targeted (Tg) to neurons by surface functionalization of the corresponding pDNA loaded NP with the nontoxic and neurotropic C-terminal 54 kDa fragment of the tetanus neurotoxin (TeNT) heavy chain (HC) [20]. Encouragingly, our previous results showcased a well-designed TMC-pDNA NP. This NP exhibits specific affinity for neurons [21]. Upon internalization, undergoes retrograde transport [16] enabling the possibility to be peripherally administered via a minimally invasive intramuscular injection [18].

Here, we proposed to deliver siRNA for downregulating PTEN expression (siPTEN) in neurons mediated by TMCSH-based NPs functionalized with the HC fragment, to enhance the vector neuron specificity. We explored the physicochemical properties of our Tg polyplexes to improve their therapeutic potential in the context of both PNS and CNS, employing various neuronal models of increasing levels of complexity. Specifically, cell lines, 3D *ex vivo* models, and neuronal cultures conducted in microfluidic systems under static and flow conditions to mimic the nervous system anatomy. Microfluidics, crucial for bridging the gap between *in vitro* and *in vivo* models while minimizing animal testing reliance [16, 22, 23], have allowed for the creation of compartmentalized neuronal cultures. This innovative approach separates neuronal cell bodies from axonal terminals, providing precise spatial control for specific, compartmentalized treatments to both axonal terminals and cell bodies. Moreover, the incorporation of flow allows the mimicking of the physiological environment, enabling the unraveling of new insights into the behavior of NPs. To the best of our knowledge, this is the first report examining the interaction of polymeric NPs in both PNS or CNS microfluidic cultures, under both static and flow conditions.

## 2. Results and Discussion

### 2.1. Development and physicochemical characterization of neuron-targeted polyplexes for siRNA delivery

Cationic polymers like chitosan can self-assemble with anionic siRNAs through electrostatic interactions forming polyplexes. Here, the formation of complexes between the TMCSH and a therapeutic siRNA was investigated (siRNA against PTEN). To follow siRNA downregulation, we have also explored the delivery of siRNA against the green fluorescent protein (GFP) in GFP expressing neurons. To confer neuron-specific targeting to the NPs, the polyplexes were functionalized with the non-toxic and neurotropic TeNT HC domain. Particularly, our Tg polyplexes were engineered to replicate the TeNT cell entry mechanism, allowing them to use the active retrograde axonal transport machinery and reach the neuron cell bodies [16, 24].

The complexation capacity and interaction strength between TMCSH and siRNA (both siPTEN and siGFP, see Table S1 in Support Information (SI) for details on the used sequences) after the tethering of the HC moieties were assessed by means of the SYBR™ Gold intercalation assay and polyacrylamide gel retardation electrophoresis (PAGE), as a function of the relation of quaternized amine groups (N) to moles of siRNA phosphate groups (P) (N/P = 2, 4 and 6; Figure 1A and Table S2). Non-targeted (nTg) polyplexes were used as control (Table S2, SI).

**Figure 1.**
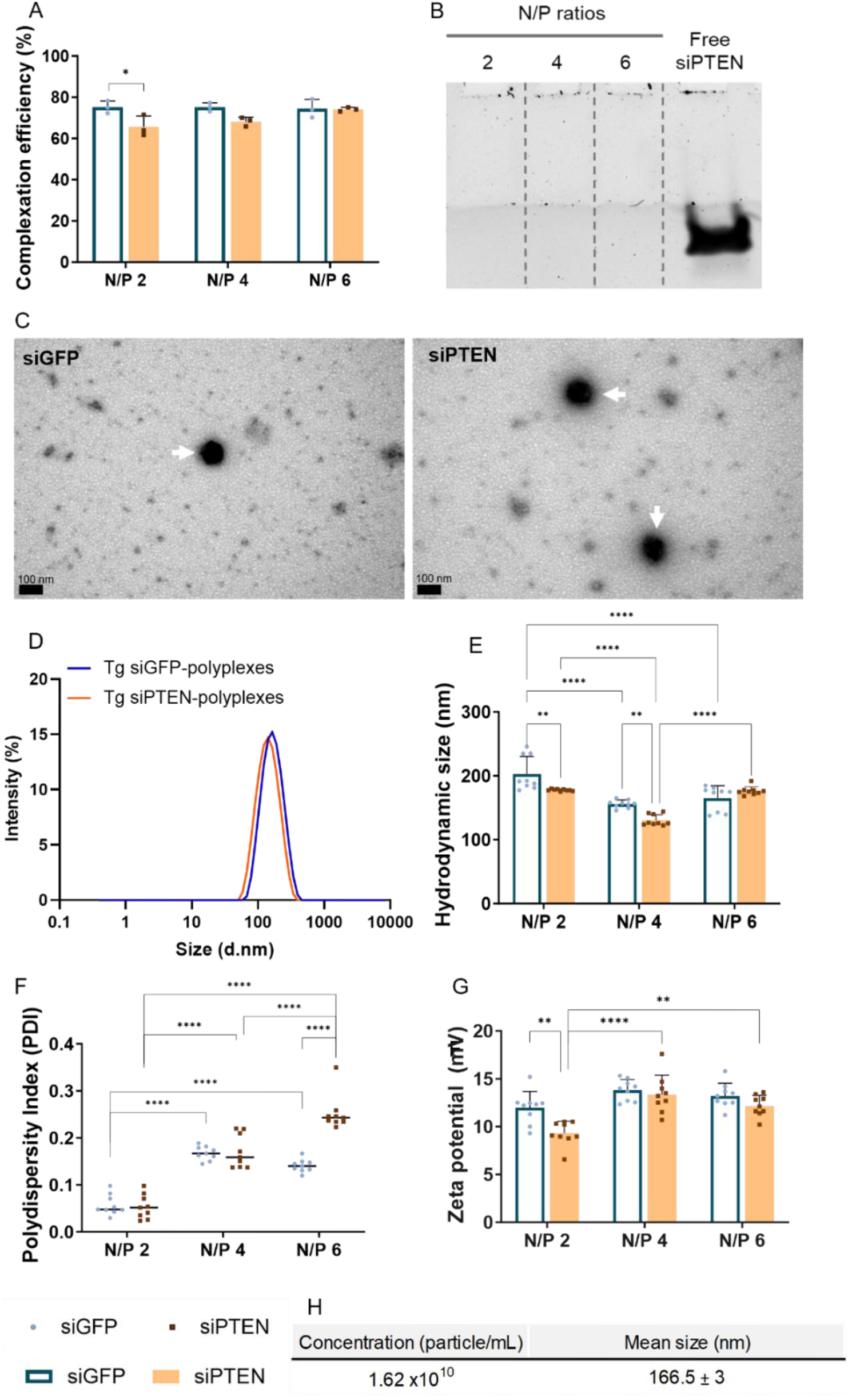
Physicochemical characterization of targeted (Tg) polyplexes prepared with siGFP or siPTEN as a function of N/P ratio. All NPs were prepared in phosphate-buffered saline (PBS) 1×, pH 7.4. (A) SYBR Gold exclusion assay (room temperature); (B) Polyacrylamide gel electrophoresis (PAGE); (C) Transmission electron microscopy (TEM) images of Tg N/P 4 polyplexes prepared with siGFP (on the left) or with siPTEN (on the right). White arrows indicate polyplexes. Scale bar: 100 nm. (D) Representative dynamic light scattering (DLS) spectrum of Tg polyplexes carrying siGFP or siPTEN; (E) Size distribution (DLS); (F) Polydispersity index (PDI, DLS); (G) Surface charge (Electrophoretic Light Scattering); (H) Concentration and mean size of Tg N/P 4 polyplexes in solution (nanoparticle tracking analysis, NTA). Results are shown as mean ± SD (standard deviation) of three independent experiments (n = 3) with (A) two or (B-H) one replicates per experiment. For statistical analysis, two-way ANOVA test was used. Significant differences: *p < 0.05, **p < 0.01, and ****p ≤ 0.0001. Results of the physicochemical characterization of non-targeted (nTg) polyplexes can be found in Table S2, Support Information.

The use of the fluorescent cationic molecule SYBR™ Gold, which selectively binds to free nucleic acids, allowed for the quantification of unbounded siRNA levels, as nucleic acids complexed with the TMCSH are inaccessible to the dye. The complexation of siGFP proved to be effective, achieving complexation rates of 64-70% and approximately 75% for both nTg and Tg formulations, respectively. Similarly effective was the complexation of siPTEN, ranging between 62-72% and 67-74% for nTg and Tg formulations, respectively. Statistically significant differences were only observed for the complexation efficiency of the Tg particles prepared with the lowest N/P ratio tested between siGFP and siPTEN (Figure 1A). The higher capacity to form complexes with siGFP, particularly relevant at lower N/P ratios where there is a minor excess of polymer, may be attributed to the distinct nucleotide sequences, namely in terms of nucleotide chemical modifications, hydrophobicity, and number (Table S1, SI). Through PAGE, it was possible to assess the migration of siRNA, which occurs only when it is loosely complexed or in free form. When Tg polyplexes were loaded into the gel and subjected to an electric field, no bands corresponding to free siRNA were identified across all tested N/P ratios (Figure 1B). Signal quantification confirmed the absence of any bands in the tested formulations (Figure S1, SI), corroborating the good complexation results obtained in the SYBR™ Gold assay. These results emphasize the vector’s siRNA complexation capacity. TMCSH has previously demonstrated great interaction capacity with siRNA. Varkouhi et al. (2010) demonstrated that TMCSH was capable of complexing with siRNA (N/P ratio of 8), causing the nucleic acid to remain in the starting plots, unable to electrophoretically migrate along the agarose gel [25]. TMCSH has also demonstrated the ability to complex other nucleic acids, namely pDNA (complexation above 90%) [17, 19]. However, it should be mentioned that the TMCSH-based NPs carrying pDNA were prepared at an N/P ratio of 15. In contrast to the studies mentioned above, the NPs in this work were prepared at N/P ratios of 2, 4, and 6, indicating a substantially lower amount of biomaterial in the NP composition. Nevertheless, the complexation remained effective, and the use of a smaller quantity of TMCSH is desirable. Considering our intention to apply these NPs to neuronal cells, formulations with reduced amount of synthetic materials are preferred [16].

The Tg and nTg polyplexes’ morphology, hydrodynamic size, polydispersity index (PDI), zeta potential, and concentration were assessed through transmission electron microscopy (TEM), dynamic light scattering (DLS), respectively (Figure 1C-G and Table S2, SI). TEM images showed that the developed Tg polyplexes, both prepared with siGFP or siPTEN, had similar spherical morphologies (Figure 1C). Across the tested N/P ratios, both Tg and nTg polyplexes exhibited narrow average sizes below 200 nm (Figure 1D and E, and Table S2, SI) and PDIs below 0.25 (Figure 1F and Table S2, SI), which is within the range of common PDI values obtained for chitosan-based NPs [26]. In general, PDIs significantly increase with the increasing N/P ratio. This could be attributed to the presence of a higher quantity of TMC, which would lead to a higher heterogeneity of the NPs. Nonetheless, both the average sizes and PDIs of Tg and nTg siRNA-polyplexes were found to be lower than those previously recorded by us with Tg and nTg pDNA-polyplexes [17]. Furthermore, both the nanosize and PDI observed here were lower than other recorded for chitosan-based NPs prepared with siRNA (sizes equal to or greater than 200 nm and PDIs above 0.3) [27]. In terms of zeta potential, both Tg and nTg formulations showed positive zeta potential across all the tested N/P ratios and siRNAs (Figure 1F and Table S2, SI). The range of surface charge of both formulations appeared very similar. The positive charge of the NPs is crucial to prevent charge-driven interaction dependent on neuronal activity, thereby reducing attraction due to neuronal spiking [28]. However, a high cationic character of the NP is also undesirable, as the interaction with cells may interfere with the receptor interactions and increase cytotoxic due to destabilization of cell membrane [29]. Thus, considering the obtained results of zeta potential (positive values but not excessive), the interaction of the Tg NPs should occur through receptors on the cellular membrane, such as TeNT receptors. In fact, the zeta potential values obtained here are very similar to those reported by Tang et al. (2022), who developed chitosan NPs modified with a targeting ligand (α-cyclam-p-toluic acid) as a novel siRNA carrier for specific delivery to injured kidneys [30]. Taken together, these results indicate that the functionalization with HC did not cause significant changes in the complexation efficiency, size, PDI, and zeta potential of the NPs. Due to slightly enhanced physicochemical characteristics, particularly in terms of size and PDI, N/P 4 was chosen as the preferred ratio to proceed with the study.

The evaluation of Tg N/P 4 polyplexes carrying siPTEN was further conducted using nanoparticle tracking analysis (NTA). The particle concentration in solution was found to be approximately 1.62 × 10^10^ particles per mL (Figure 1H). NTA was also used to determine the mean particle size in suspension. Tg polyplexes were found to present sizes of 167 ± 3 nm, closely matching those obtained by DLS. The stability of Tg N/P 4 polyplexes in the cell culture medium used for cell lines, dorsal root ganglion (DRG), and primary neuronal cultures was also verified by DLS (Figures S2, SI). The results demonstrated minimal variation in the hydrodynamic diameter of the polyplexes across different media, indicating that Tg N/P 4 polyplexes maintained their structural integrity. This stability is crucial for ensuring consistent performance in cellular uptake and gene delivery, as significant size changes could affect their interaction with cells and overall therapeutic efficacy.

### 2.2. Enhanced cellular uptake and neurospecificity of targeted polyplexes

Firstly, the cellular viability of neuronal cells following 24 hours of exposure to Tg and nTg polyplexes (N/P 4) carrying siPTEN was assessed. A lactate dehydrogenase (LDH) cytotoxicity assay was performed, as well as the measurement of spontaneous neuronal activity. To the best of our knowledge, this is one of the first reports showing the impact of the treatment of neuronal cell populations with NPs on spontaneous neuronal activity.

Cultures of mice embryonic primary motor and cortical neurons were established under standard conditions. Both Tg and nTg NPs were found to cause minimal impact on the cell membrane. This conclusion is based on the low percentage of released LDH observed in the motor (Figure S3A, SI) and cortical neurons (Figure S3B, SI) after exposure to these NPs. Noteworthy, significant differences were found in terms of LDH release when both PNS and CNS neurons were treated with the commercially available lipid vector -lipofectamine. Lipofectamine was used as a control since it is considered one of the standards for in vitro transfections.

For nucleic acid delivery vectors to be biologically applicable in the context of the nervous system, vectors should not damage the electrophysiological response of neuronal cells, as this could indicate a lack of biocompatibility and interference with neuronal communication. The impact of Tg and nTg TMCSH-based NPs carrying siPTEN on neurons involved the analysis of their influence on cells’ electrical activity (Figure S4, SI). The formulations’ effects were evaluated in neurons cultured in commercial six-well-plate microelectrode arrays (MEAs) (nine electrodes per well), after exposing the entire cell population to the NPs. Following the measurement and analysis of motor neuron activity (Figure S4A-F, SI), cells treated with Tg or nTg polyplexes displayed spontaneous electrical features (number of spikes, number of bursts, interspike interval (ISI), and mean intra-burst interval (IBI)) similar to untreated cells. Likewise, the activity of cortical neurons remained the same after incubation with Tg and nTg polyplexes, closely matching the electrical activity values recorded in untreated cell cultures (Figure S4G-L, SI). Furthermore, the neuronal spiking data was similar between NPs treated and untreated samples (Figure S4M, SI). Most electrodes were active, showing a heterogeneous firing-rate distribution (Figure S4N and O, SI). This indicates that the neuronal population in the culture plate underwent no electrophysiological alterations upon interaction with NPs. As expected, the neuronal electrical activity was only altered when treated with Triton X-100. This electrophysiological response further reinforces the neuro-applicability of the developed vectors.

As the biocompatibility of our vectors towards sensitive cells, such as primary mouse neuronal cells, was assured, the specific binding capacity of the NPs was investigated. For this, neuronal cell lines (ND7/23 and HT22, mouse neuroblastoma and rat neuron hybrid and mouse hippocampal neuronal cell lines, respectively) were chosen due to the known presence of GT1b receptors in their membrane composition [31, 32]. These receptors are recognized and have affinity for the binding domain of the TeNT. A non-neuronal cell line (NIH 3T3, mouse embryonic fibroblast cell line) was used as the negative control. Initially, through flow cytometry, the binding capacity of nTg polyplexes to different neuronal and non-neuronal cell types was assessed (Figure S5, SI). nTg polyplexes at N/P ratios 2, 4 and 6 were prepared with a mimetic of siGFP (siRNAmi) labelled at the 5′ end of the sense strand with cyanine 5 (Cy5-siRNAmi) and added to the cells. After 24 hours of incubation, the Cy5 signal was observed in all cells in all conditions (nearly 100 % Cy5-positive cells) (Figure S5, SI). With the increase in the N/P ratio, there was an increase in the signal intensity associated with the cells, suggesting that the presence of a higher amount of TMCSH in the complex formulation allows for more efficient transport of the Cy5-siRNAmi to the cells. Overall, the nTg polyplexes demonstrated a good capacity for interaction with cells, independent of their origin.

Subsequently, the binding capacity of HC-functionalized polyplexes to both neuronal and non-neuronal cells was evaluated (Figure 2A). Higher percentages of Cy5-positive cells were observed with the increase of the N/P ratio. Between 85 and 99% of ND7/23 cells showed positive Cy5 signal, and similarly high binding was observed with HT22 cells, where there were between 79 and 95% positive cells. Contrariwise, the binding to NIH 3T3 cells was significantly lower, with values between 50 and 60% of positive cells for Cy5 signals. The clearly higher percentage of Cy5 signal observed in the neuronal cell lines was expected due to the functionalization with the TeNT binding domain. The HC enhanced the binding of Tg polyplexes to ND7/23 and HT22 cells, indicating a greater propensity of neuron-Tg NPs to interact with neuronal cell lines. Regardless of the cell line, lipofectamine exhibited a high percentage of cells with Cy5 signal (above 90%). However, this indicates that internalization was not specific, as all three cell lines showed statistically identical results. When the pre-incubation of the cells with free HC was performed before the treatment with the Tg N/P 4 polyplexes, a much lower intensity of Cy5 associated with neuronal cells was noted, indicating reduced binding of the NPs to these cells (Figure 2B). Only 5-12% of ND7/23 cells exhibited Cy5 fluorescence signal and merely 7-14% in HT22 cells. This suggests that the free HC has blocked the receptors with which the Tg NPs interact. Conversely, pre-incubation with the TeNT fragment did not induce alterations in the interaction with NIH 3T3 cells (56-77% Cy5-positive cells). Since the percentage of positive cells remained unchanged with pre-treatment using HC, the entry of the polyplexes into these non-neuronal cells is thus considered non-specific. Altogether, the association of Tg polyplexes was preferential and occurred to a greater extent with neuronal cells. The superior results obtained with the NPs prepared with an N/P ratio of 4, indicating better overall performance in terms of physicochemical characteristics and good binding capacity to neuronal cells, reinforced our decision to use these particles to continue the study.

**Figure 2.**
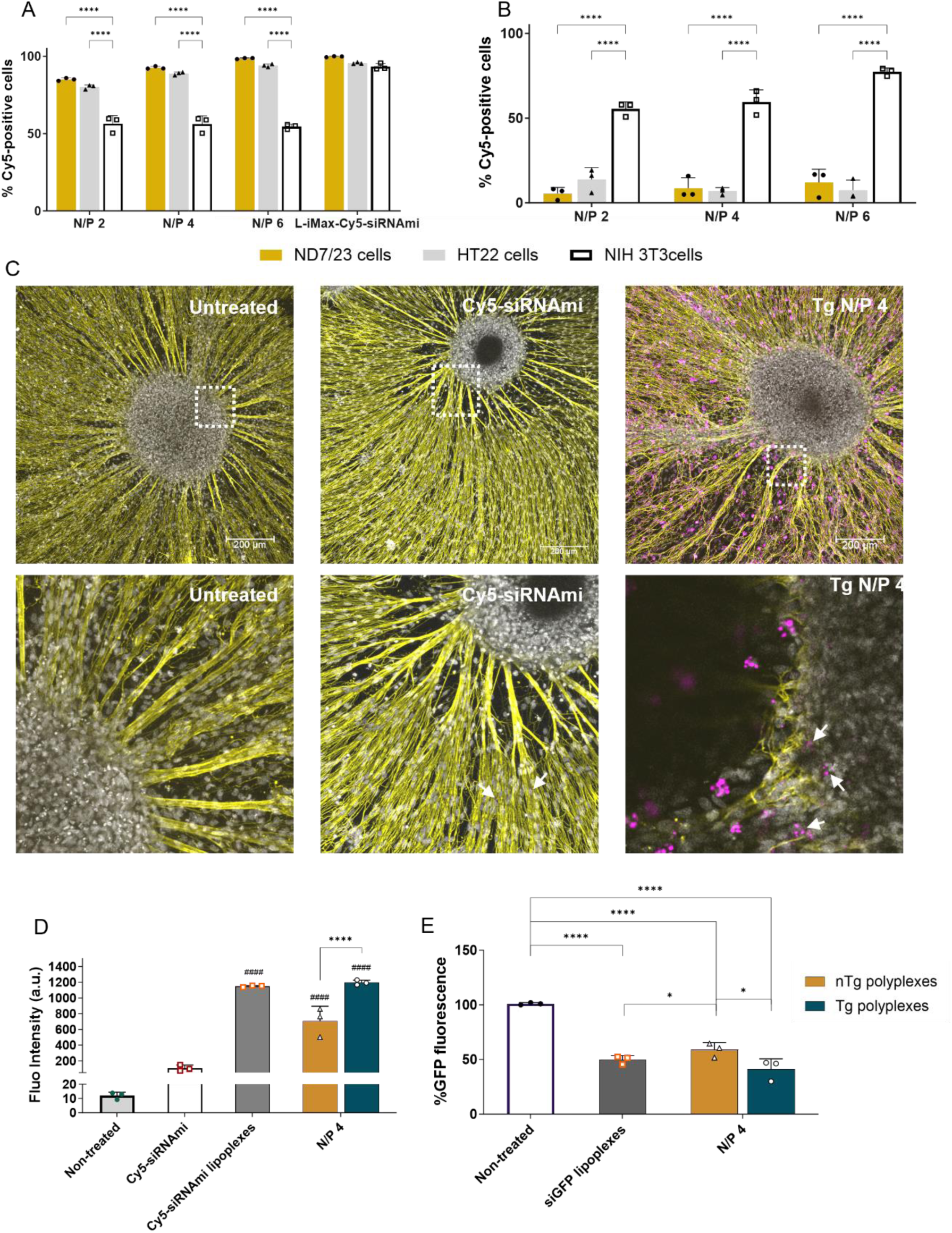
Cellular binding, internalization and downregulation capacity of polyplexes. (A) ND7/23, HT22 and NIH 3T3 cells were incubated with targeted (Tg) polyplexes (N/P 2, 4 and 6) carrying Cy5-labeled siRNAmi for 8 hours (final siRNAmi concentration 100 nM). Characterization by flow cytometry in terms of Cy5 positive cells (in %); (B) A similar cellular association assay was done but including a pre-incubation with HC (final concentration 1 nM) for 15 minutes at 4 ͦC; (C) Representative images of entire embryonic DRG explant untreated (on the left) or incubated, for 24 hours, with Cy5-siRNAmi (on the center) and with Tg polyplexes carrying Cy5-siRNAmi (final concentration 100 nM, on the right). Staining: βIII-tubulin (yellow), nuclei with Hoechst 33 342 (gray), and Cy5-siRNAmi (pink). White arrows indicate Cy5 signal associated to the polyplexes in the DRG explant body Scale bar: 200 μm; (D) Cy5 fluorescence intensity quantification in the DRG explant body. (E) Downregulation of GFP expression measured in motor neurons after 96-hour incubation with N/P 4 polyplexes. Results are represented as mean ± SD of three independent experiments (n = 3), with two replicates per experiment. For statistical analysis, two-way ANOVA test was used. Significant differences: *p < 0.05, and ****p ≤ 0.0001.

The internalization capacity of both Tg and nTg N/P 4 polyplexes was subsequently in cultured DRG explants using confocal microscopy (Figure 2C and D). Both polyplexes, complexing Cy5-siRNAmi, demonstrated remarkable internalization capability within this more complex 3D model. Their internalization was observed in both the core of the DRG, *i.e*., the neuronal cell bodies, and in satellite cells (Figure 2C). Furthermore, it is noteworthy that DRG cultures did not show any visual signs of compromised viability. Signal quantification revealed that DRGs treated with Tg polyplexes exhibited a significantly higher accumulation of Cy5 signal than DRG treated with free Cy5-siRNAmi or nTg polyplexes (Figure 2C and D). Quantitatively, DRGs treated with Tg polyplexes exhibited a 1.7 times higher Cy5 fluorescence signal in the explant body compared to those treated with nTg polyplexes. Moreover, the fluorescence intensity of DRGs treated with Tg polyplexes (1196 ± 32 a.u.) was very close to the intensity of DRGs treated with the lipidic standard - lipofectamine (around 1149 ± 11 a.u.). This result is highly promising, demonstrating that our vector also possesses a high internalization capacity, namely in complex models such as the 3D cultured DRG explant.

Considering the enhanced neurospecific association showed, the downregulation effect on a reporter gene (GFP) was evaluated (Figure 2E). Primary motor neurons expressing the HB9-eGFP fusion protein were treated with Tg or nTg (N/P 4) carrying siGFP (final concentration of 100 nM) for a 96-hour incubation period. The transfection efficiency was determined by the reduction in GFP reporter gene expression quantified through image analysis. Treatment with both types of NP formulations induced a decrease in fluorescence intensity. Non-targeted polyplexes led to a 51% reduction in GFP expression, while Tg polyplexes achieved a significantly greater reduction, silencing approximately 59% of the expression (Figure 2E). The functionalization with the TeNT HC domain is expected to have contributed to this enhanced transfection efficiency. Furthermore, the downregulation effect of Tg polyplexes closely resembled the outcome observed in cells treated with the standard lipidic vector, which resulted in a 50% reduction in GFP expression. Despite the substantial silencing achieved by lipofectamine, its *in vivo* applicability is limited due to toxicity. In contrast, our polyplexes emerge as a promising option, supported by demonstrated biocompatibility (previously in this section) and comparable transfection efficiency.

#### The interaction of targeted polyplexes with axons under flow conditions

The binding of Tg and nTg N/P 4 polyplexes, transporting Cy5-siRNAmi, was then assessed in dynamic microfluidic cultures of neurons (either motor or cortical), which effectively isolate cell bodies from axonal terminals (Figure 3) in a more physiological manner [16]. A significant difference from the other models used in this study is the exploration of a peripheric administration (only in the axonal compartment of a two-compartment microfluidic device) and the inclusion of flow conditions (Figure 4A and B). The study of the binding efficiency of NPs to other types of cells using *in vitro* flow adhesion assays has been previously described [33, 34], and proved that flow conditions may alter the interaction of NPs with cell populations. As far as we know, this study represents a pioneering use of microfluidic systems with flow to assess the neuron-specificity of NPs.

**Figure 3.**
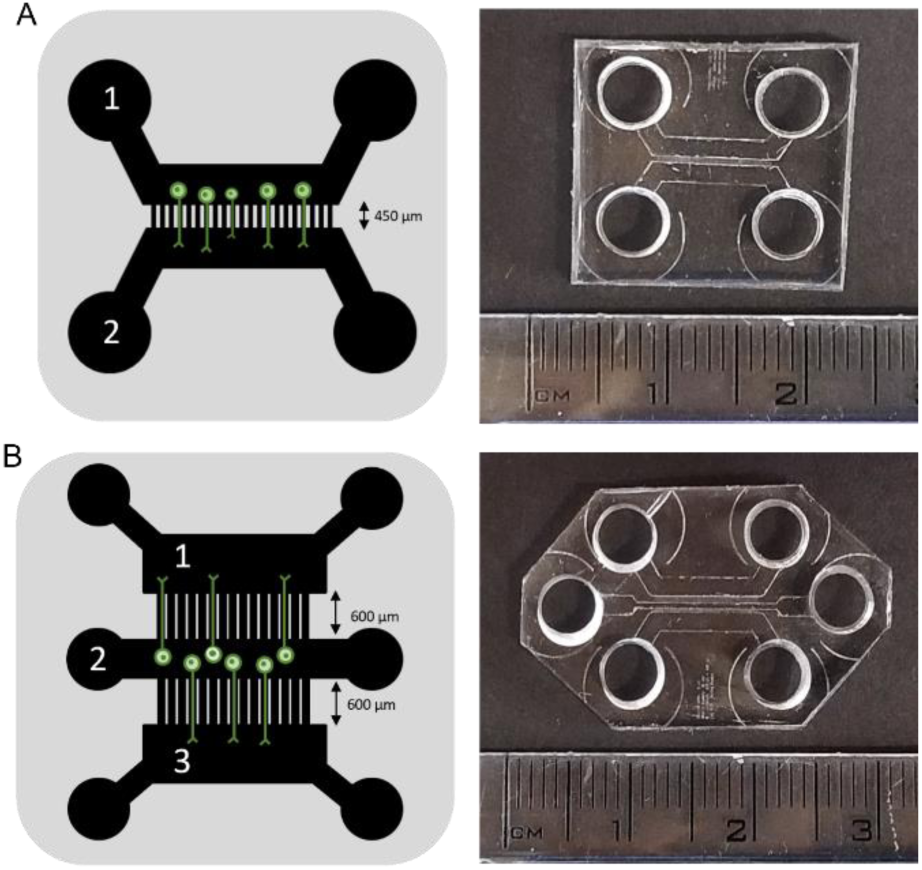
Schematic representation of the microfluidic chips design and pictures of the poly(dimethylsiloxane) (PDMS) microfluidic devices employed in this study. (A) Two-compartment microfluidics with 450 µm microchannels. Compartment 1 refers to the cell soma compartment, while compartment 2 is the compartment for neuronal axon terminals; (B) Three-compartment microfluidics with two sets of 600 µm microchannels. Compartments 1 and 3 refer to the compartments for neuronal axon terminals, and compartment 2 is the cell soma compartment of the neurons.

**Figure 4.**
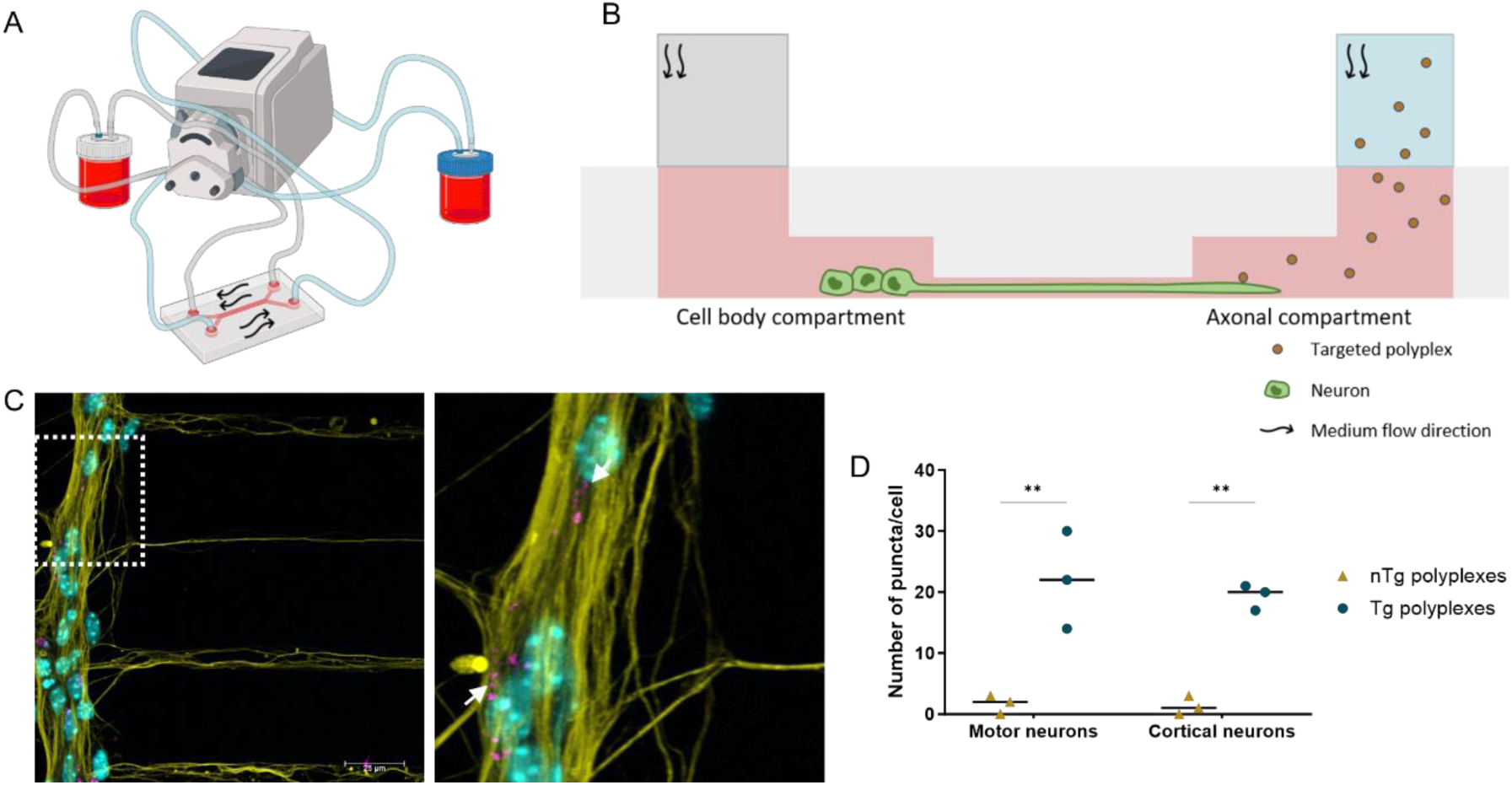
Polyplexes interaction with neurons, under flow conditions. Motor or cortical neurons were seeded in two-compartment microfluidics and incubated overnight under flow with Tg and nTg N/P 4 polyplexes carrying Cy5-labeled siRNAmi (added in axonal compartment, final concentration 100 nM). (A) Schematic representation of the setup; (B) Side-view illustration of nanoparticle incubation under flow in microfluidic; (C) Representative image of cortical neurons incubated with Tg N/P 4 polyplexes carrying Cy5-siRNAmi. Staining: βIII-tubulin (yellow), nuclei with Hoechst 33 342 (cyan), and Cy5-siRNAmi (pink). White arrows indicate Cy5 signal associated to the polyplexes. Scale bar: 25 μm; (D) The number of puncta inside neurons (cell soma compartment) was quantified. Results are represented as mean ± SD of three independent experiments (n = 3), with one replicate per experiment. For statistical analysis, two-way ANOVA test was used. Significant differences: **p < 0.01.

The NPs were administered in a constant circulating culture medium, passing through the compartment containing only axonal terminals. Twenty-four hours after the addition of polyplexes, the cells were fixed and analysed using a confocal microscope (Figure 4C). Interestingly, neurons incubated with Tg polyplexes under flow exhibited significantly more puncta with the fluorescence signal of Cy5-siRNAmi (Figure 4D). The behaviour of Tg NPs with both types of neurons (motor and cortical neurons) appears similar, what can be expected as TeNT has demonstrated the ability to interact with both PNS and CNS neurons [35, 36]. On the other hand, neurons treated with nTg polyplexes did not show any considerable signal corresponding to the presence of nucleic acid. These results emphasize the importance of the use of a specific targeting moiety in NPs’ composition. Targeting moieties are essential for specifying a treatment to a particular cell type or tissue, reducing side effects. Importantly, under flow conditions, these molecules were found crucial for enabling particles to interact with neurons. Under dynamic conditions, for interactions to be effective, the bonds should be stronger, a feat achieved only through ligand-receptor interactions. The superior strength of the interaction between HC and receptors was previously determined by us using atomic force microscopy [21]. In a previous study we have shown that higher forces are required to break the bond between HC-functionalized NPs and the membrane of DRG neurons and neuronal tissues. Thus, the incorporation of specific targeting moieties not only enhances the selectivity and efficacy of NPs interactions with neuronal cells but also underscores the necessity of strong ligand-receptor bonds to assure these interactions under dynamic conditions.

In summary, the TeNT HC fragment proved to be important in enhancing the preferential and effective interaction of Tg polyplexes with neuronal cells.

### 2.3. Efficient axonal retrograde transport of neuron-targeted polyplexes

Two-compartment microfluidic chips (Figure 3A) were further utilized as a model to evaluate if the tethering of the HC fragment to the polyplexes improved the retrograde transport of the NPs toward the cell body of neurons. This model allows the evaluation of NP uptake at axon terminals and subsequent possible axonal transport to the cell body of the cell [16]. To explore this, live cell imaging experiments were conducted using motor or cortical neurons cultured within the microfluidic devices. Polyplexes were introduced into the medium of the axonal terminals’ compartment, and subsequent NP transport was visualized in the axons growing within the microchannels. Live imaging analysis revealed the retrograde transport of numerous NP-loaded vesicles along axons after internalization in their terminals at the axonal compartment (Figure 5 and S6, SI). Through the tracking of NP trafficking along the axons, the motion characteristics of loaded vesicles for both Tg and nTg polyplexes were analysed using various parameters. These included the probability density function of instantaneous velocity, average velocity and time spent in movement versus pause. The probability density function of instantaneous velocity, which is a function that describes the continuous velocity of particles in motion, exhibited greater variability and a higher median instantaneous speed for Tg polyplexes-loaded vesicles in both motor and cortical neurons (Figure 5A and B). In the nTg polyplex treated cells a more uniform velocity distribution was determined, but significantly lower (between −0.3 and 1 µm/s). The distribution of instantaneous speeds in motor and cortical neurons were found quite similar for both nTg and Tg polyplexes. Interestingly, in both motor and cortical neurons vesicles containing nTg NPs exhibited more anterograde movements, characterized by negative values of instantaneous speed, indicating short reversals in direction for vesicles moving predominantly in the retrograde direction (Figure 5A and B). The distribution of instantaneous speeds and movement direction observed here is reminiscent of our previous findings with TMCSH-based NPs carrying pDNA, both with Tg and nTg formulations, evaluated in DRG neurons [16]. Even though movement-related properties were studied in three different types of neurons (motor, cortical, and DRG neurons), these properties remained consistent.

**Figure 5.**
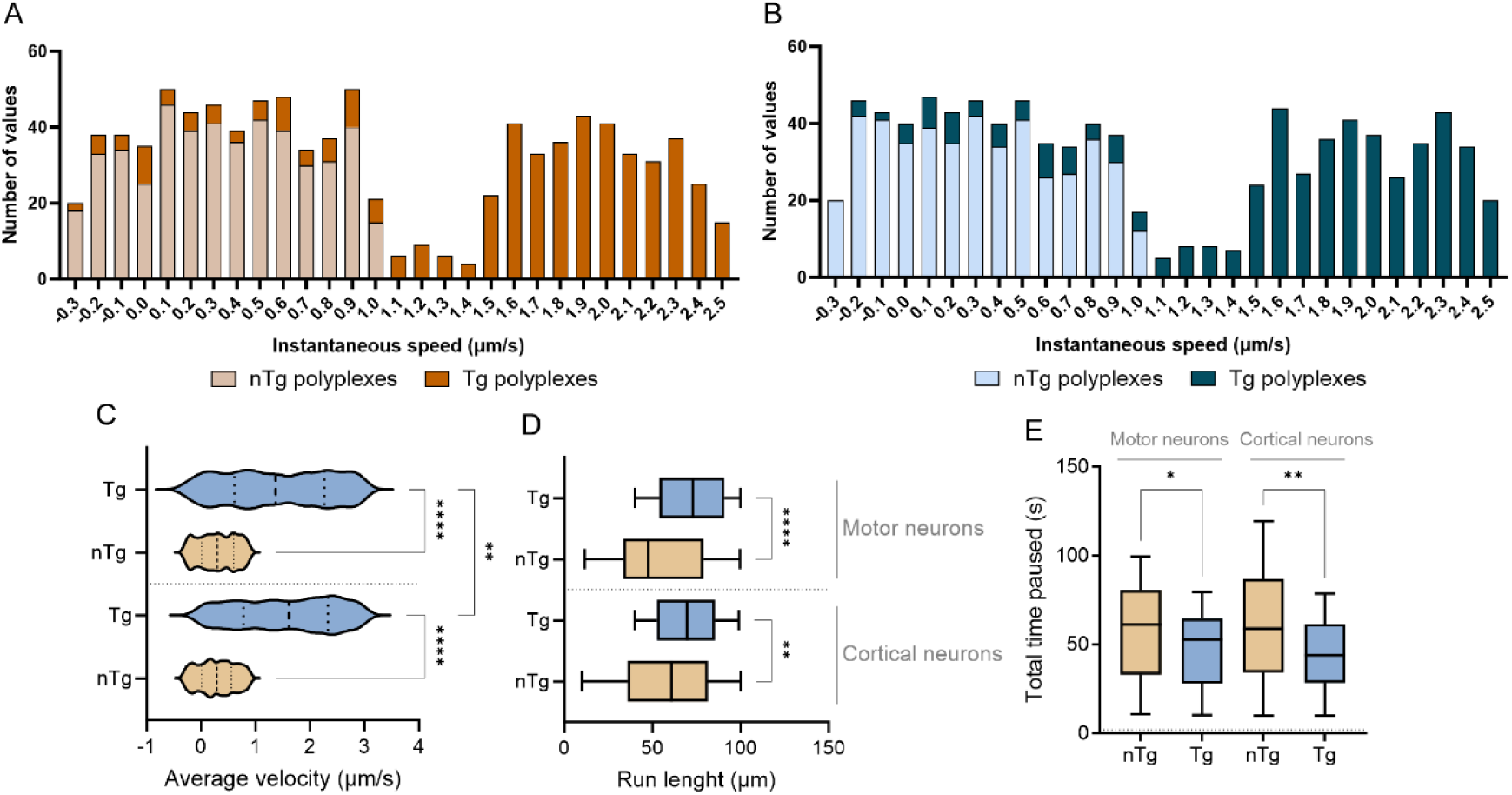
Retrograde axonal transport characteristics in motor and cortical neurons. Probability density function of the instantaneous velocity of nTg and Tg N/P 4 polyplexes-loaded vesicles in (A) motor neurons and (B) cortical neurons, defined as the speed (μm/s) of movement between consecutive images within a sequence; (C) Average velocity (μm/s) of retrograde axonal transport of NPs-loaded vescles; (D) total run length and (E) total pause periods. Results are represented as mean ± SD of three independent experiments (n = 3), with one replicate per experiment. One-way ANOVA tests were used for statistical analysis. Significant differences: *p < 0.05,**p < 0.01, and ****p ≤ 0.0001.

Furthermore, when examining average velocity values, both in motor and cortical neurons, vesicles loaded with Tg formulations demonstrated transportation at high velocities (superior to 1 μm/s), falling within the range of fast axonal transport (Figure 5C) [37]. In motor neurons, the average velocity of Tg polyplexes was around 1.40 µm/s, while in cortical neurons, the average velocity was approximately 1.55 µm/s, both in the range of fast axonal transport. On the other hand, the average velocity of the nTg NPs was significantly lower, around 0.3 μm/s in both neuronal primary cells. Therefore, the average velocity of Tg polyplexes was 4.7 times higher in motor neurons, and in cortical neurons, the average velocity was 5.2 times greater. This observation underscores that surface functionalization with the HC fragment enhances both the retrograde axonal transport capacity and migratory velocity of TMCSH-based NPs. It is key to note that the average velocities of both Tg and nTg polyplexes closely match the velocities of the microtubule-dependent molecular motor dynein, responsible for retrograde cargo transport [37, 38].

Regarding the total distance moved per run, the Tg polyplexes exhibited a significantly greater coverage (Figure 5D). On average, Tg polyplexes migrated 68.8 µm along motor neuron axons and 72.4 µm in cortical neurons. The average distance covered by Tg systems was 1.2 times greater in motor neurons, and 1.4 times greater in cortical neurons compared to nTg systems. While both types of neurons showed considerable coverage by nTg polyplexes, the HC conferred an enhanced ability to the NPs to cover even longer distances.

HC functionalization also leads to a significant reduction of the time during which the NPs are stationary. Assessing the overall duration of NP pausing (no movement along axons), the Tg polyplexes showed less stationary time compared to the nTg polyplexes in motor neurons (Figure 5E). Overall, the functionalization with HC endowed the polyplexes with the ability to move with fewer interruptions, maybe due to more frequent interactions with microtubule motors. For instance, the dynein motor undergoes short-range movements of 1-2 μm in the microtubule, implying that long-range transport is achieved through the continuous exchange of motile dynein complexes [39, 40]. These exchanges can thus justify the recorded pauses seen in Tg polypexes movement. Curiously, significant differences in the run length and total time paused had not been previously observed by us when assessing these parameters for Tg and nTg pDNA-polyplexes [16]. The absence of changes in the distance migrated and time paused may be attributed to the properties of the NPs, namely the smaller hydrodynamic sizes of the siRNA-polyplexes. However, it could also be a result of the distinct neuronal type, which may exhibit different axonal transport mechanisms.

Furthermore, to assess the potential of neuron-Tg polyplexes in effectively reaching the cell body of neurons, nTg or Tg polyplexes, prepared at N/P ratio 4 and carrying Cy5-siRNAmi, were introduced into the axonal terminals compartment for 12, 24, or 48 hours. At the designated time points, the cells were fixed and subsequently analysed through confocal microscopy (Figure 6). The Cy5 fluorescence signal was quantified in the cell soma of the neurons (Figure 6A). Both motor and cortical neurons exhibited detectable Cy5 fluorescence after 12 hours of incubation with both types of TMCSH-based polyplexes (Figure 6B and C). As the incubation time increased, the fluorescence signal gradually and continuously intensified both in motor and cortical neurons, reaching its peak for the cell incubated for 24 hours. At this time point, the cell bodies of both motor and cortical neurons displayed a 3.0-fold greater intensity of Cy5 fluorescence, compared to neurons that were incubated with nTg polyplexes. Overall, Tg polyplexes demonstrated a stronger propensity for reaching and accumulating in the neuronal cell body, especially at 12 and 24 hours of incubation (Figure 6B and C).

**Figure 6.**
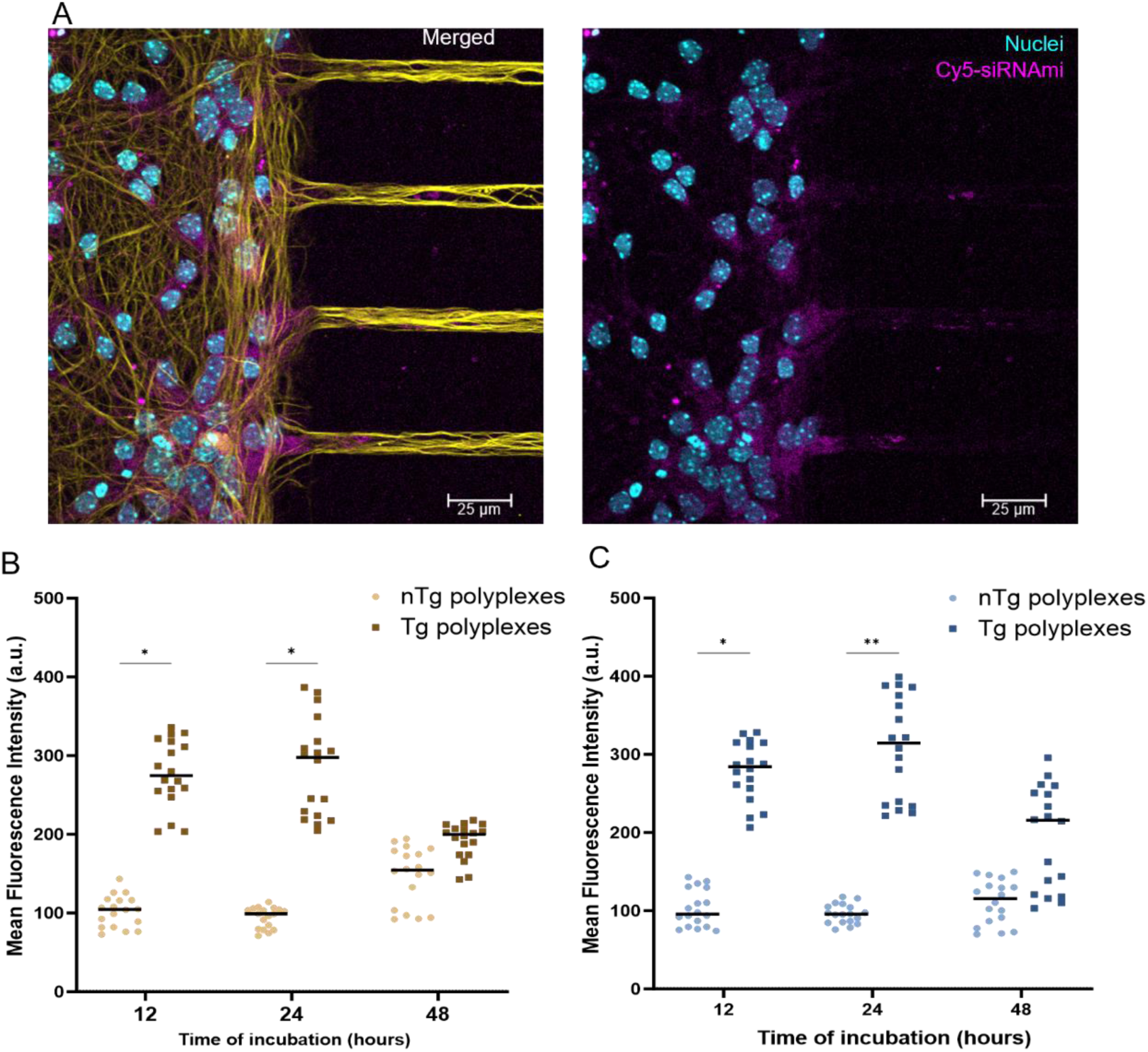
The cellular interaction kinetics of polyplexes in microfluidic-based cultures. (A) Representative images from confocal microscopy of Tg N/P 4 polyplexes carrying Cy5-siRNAmi. Staining: Nuclei with Hoechst 33 342 (cyan), βIII tubulin (yellow) and Cy5-siRNAmi polyplexes (pink). Through image analysis, the fluorescence intensity in the cell bodies of (B) motor neurons and (C) cortical neurons was quantified. Results are represented as the mean ± SD of three independent experiments (n = 3), with at least six replicates per experiment. For statistical analysis, one-way ANOVA test was used. Significant differences: *p < 0.05 and **p < 0.01.

These observations suggest that the processes of binding and internalization for each NP formulation might be distinct, potentially resulting in different efficiencies in retrograde axonal transport [16, 41, 42]. Additionally, it is noteworthy that beyond the fluorescence intensity peak at 24 hours, a decline in the Cy5 signal intensity at the 48-hour mark was identified. This reduction could potentially be ascribed to vesicle dissociation followed by siRNA degradation.

### 2.4. Biocompatibility and biological effect of siPTEN-polyplexes

To gauge cellular viability following 24 hours of exposure to Tg and nTg polyplexes (N/P 4) carrying siPTEN, an LDH cytotoxicity assay was performed using culture medium collected from cell soma compartments (Figure S7, SI).

Primary motor and cortical neuron cultures were seeded in two-compartment microfluidic conditions and incubated with Tg or nTg N/P 4 polyplexes (added in the axonal compartment). Overall, both Tg and nTg NPs demonstrated minimal impact on the cells. This assertion is drawn from the low percentage of released LDH observed in embryonic motor neurons (Figure S7A, SI) and cortical neurons (Figure S7B, SI).

In microfluidic cultures, cells treated with Tg and nTg N/P 4 polyplexes in the axonal compartment showed very low LDH release (below 11 %). Trend-wise, the level of LDH released by cells seeded in microfluidic chips appeared slightly lower than the amount released by neurons in cells seeded in multiwell plates (Figure S3, SI), after treatment with NPs (compared to untreated cells seeded in microfluidics or plates, respectively). This result can be attributed to the lower number of NPs reaching the cell soma compared to the number of NPs contacting the cell soma in multiwell plates. Interestingly, this result aligns with our previous studies of TMC-based NPs in two-compartment microfluidic digested DRG cultures [16]. The findings in the microfluidic platforms are important, as this model faithfully replicates the physiological separation between the cell soma and axonal terminals. The microfluidic model allows NPs to interact with neuronal axons in a manner closely resembling *in vivo* conditions.

To assess spontaneous neuronal electrical activity in two-compartment microfluidic cultures, the electrical activity of neurons was measured after incubation with NPs (Figures 7 and S8, SI). No electrophysiological alterations were identified in either motor neurons (Figures 7A and S8A-F, SI) or cortical neurons (Figures 7B and S8G-L, SI) following incubation with nTg or Tg N/P 4 polyplexes. The total number of spontaneous spikes and other measures of neuronal electrical activity (number of bursts, ISI, and IBI) were similar to those in the untreated group. This demonstrates that the addition of NPs to the axons did not affect neuronal viability or activity. Furthermore, the spiking data was comparable between samples treated with NPs and untreated samples (Figure S8M, SI).

**Figure 7.**
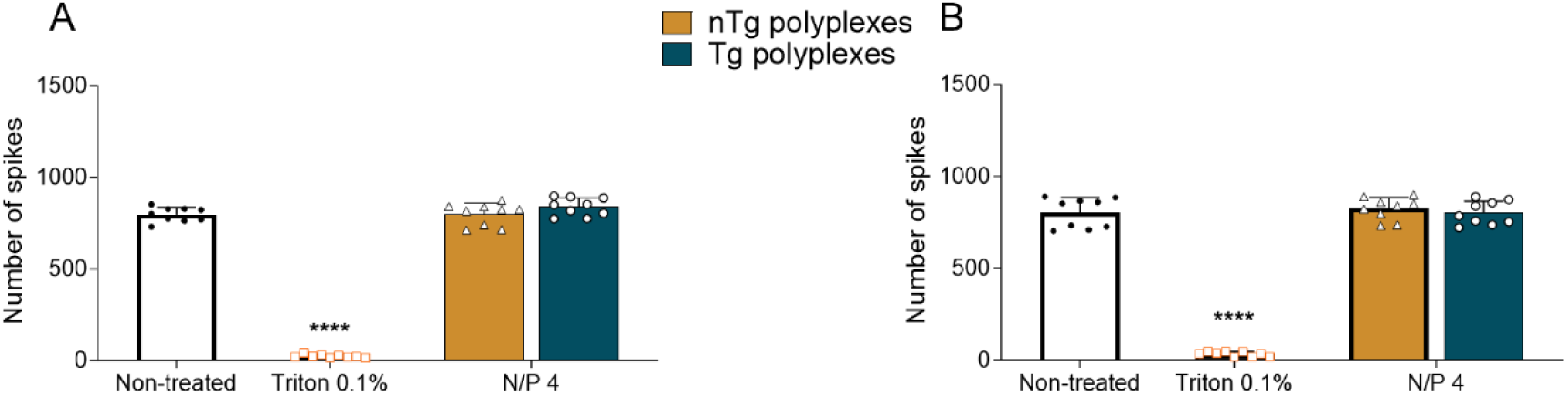
Biocompatible profile evaluation in primary motor and cortical neurons. Electrical activity evaluation through the total number of spikes of (A) motor neurons and (B) cortical neurons cultured two-compartment microfluidic. Results are represented as the mean ± SD of three independent experiments (n = 3), with three replicates per experiment. For statistical analysis, two-way ANOVA test was used. Significant differences: ****p ≤ 0.0001.

This electrophysiological response further strengthens the neuro-applicability of the developed vectors.

The biocompatible polymeric NPs, both Tg and nTg variants, underwent additional scrutiny to evaluate their ability to transport and release siPTEN. This assessment aimed to gauge their effectiveness in silencing the expression of a biologically significant gene -PTEN (Figure 8). The downregulation was indirectly assessed based on axonal outgrowth (Figure 8A), as PTEN negatively impacts this event. The experiment was conducted using the three-compartment microfluidic platforms with two sets of microchannels, each measuring 600 µm in length (Figure 3B). Primary cortical neurons were seeded in the central compartment, and within this same compartment, Tg and nTg siPTEN-polyplexes were added to scrutinize the changes in axon development resulting from PTEN silencing. Five days after adding the NPs, the cultures were fixed, and axonal growth was assessed through image analysis. Cells treated with soluble TMCSH were used as control.

**Figure 8.**
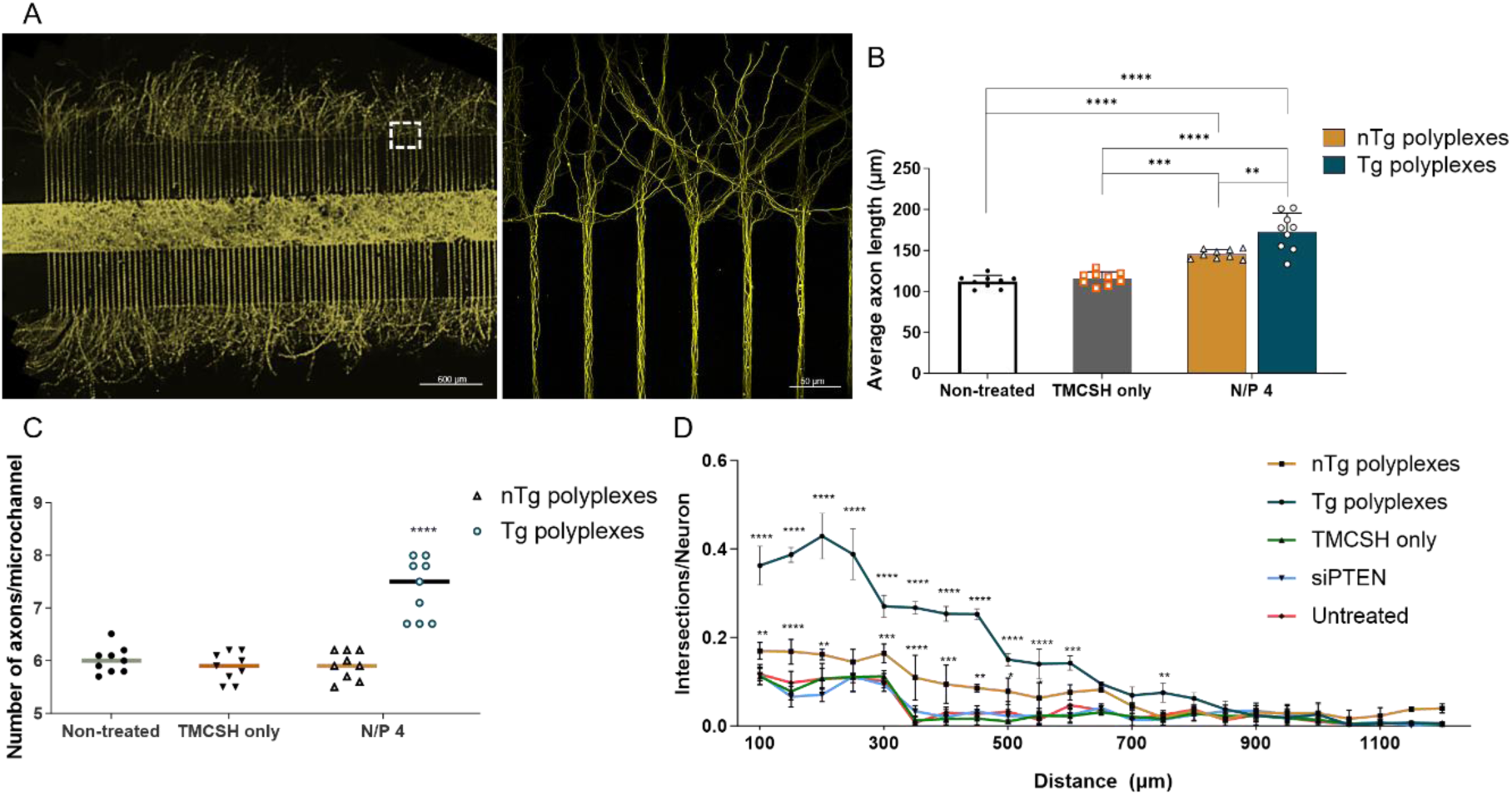
Biological effect of NPs formulations in primary cortical neurons. (A) Representative mosaic image used to quantify axonal outgrowth, in three-compartment microfluidic devices, after treatment with targeted N/P 4 polyplexes carrying siPTEN. Staining: Neurofilament H (in yellow). (B) Average axon length quantified in neurons growing in three-compartment microfluidic cultures; (C) number of axons per microchannel and (D) number of intersections of the axons in linear grids spaced 50 µm apart. Results are represented as the mean ± SD of three independent experiments (n = 3), with three replicates per experiment. For statistical analysis, two-way ANOVA test was used. Significant differences: *p < 0.05, **p < 0.01, *** p < 0.001 and ****p ≤ 0.0001. In (C) and (D), significant differences are observed compared to untreated cells.

The average length of axons that were treated with nTg and Tg polyplexes was significantly increased, measuring on average 146 and 173 µm, respectively (Figure 8B). These lengths were 1.3 and 1.5 times greater than the average axon length of untreated cells,, indicating a notable enhancement in axonal growth. Nevertheless, Tg polyplexes exhibited a more favourable axonal response compared to the other nTg formulations, (Figure 8C). Furthermore, treatment with Tg NPs led to a significantly higher number of axons that migrated within the microchannels and reached the axonal compartment, with approximately 8 axons per microchannel observed when treated with Tg polyplexes. On the other hand, the axonal response to treatment with both nTg polyplexes and the remaining controls was lower (around 6 axons per microchannel). Linear Sholl analysis MATLAB scripts were applied to analyse axonal growth within the axonal compartments (Figure 8D) [43]. When neurons were treated with nTg or Tg polyplexes, a significantly higher number of intersections per grid line were recorded, indicating that the axonal network proliferated in greater numbers and over longer distances, extending up to 900 µm.

Altogether, the HC functionalization enhanced the vectors’ therapeutical potential. The increased axonal outgrowth and boost in axon length underscore the potential of Tg polyplexes for enhancing axonal development, attributed to the effective silencing of PTEN. These findings represent a promising avenue for advancing our understanding of neural growth and regenerative therapies.

## 3. Conclusions

Our study focused on exploring and characterizing neuron-Tg TMCSH-based polyplexes for siRNA delivery, functionalized with the neurotropic TeNT HC domain as the neuron-targeting moiety.

Effective complexation of different siRNA sequences (siGFP and siPTEN) with TMCSH was demonstrated across various N/P ratios, with high percentages of complexation being obtained for both siRNAs. The polyplexes showed suitable sizes, consistently below 200 nm, with narrow PDIs and positive zeta potential. Importantly, the addition of the TeNT HC domain did not induce significant changes in NP characteristics, furthering protecting the siRNA on the NPs.

The neurospecificity of Tg polyplexes was established. Moreover, the HC-functionalized polyplexes exhibited enhanced binding to neuronal cells and superior efficiency in downregulating GFP expression in primary motor neurons, outperforming nTg polyplexes and even a commercially available transfection vector (lipofectamine). Microfluidic cultures under flow conditions further confirmed Tg polyplexes’ substantially higher uptake compared to nTg polyplexes, representing a pioneering approach in assessing neuron-specificity under flow conditions. With the TeNT fragment playing a crucial role in enhancing Tg polyplexes’ preference for neuronal interactions.

Exploring intracellular dynamics, live cell imaging experiments using a two-compartment microfluidic chip model demonstrated retrograde transport of Tg polyplexes along axons. Tg polyplexes exhibited improved motion features, higher average velocities aligning with fast axonal transport, and greater distances covered along axons compared to nTg polyplexes. HC functionalization also reduced the time polyplexes were stationary, suggesting improved interactions with microtubule motors. In reaching neuronal cell bodies, Tg polyplexes showed a stronger propensity for accumulation than nTg polyplexes.

Comprehensive biocompatibility evaluations, particularly focusing on electrical neuronal activity, confirmed that polyplexes did not induce electrophysiological alterations in neurons, supporting their compatibility with neuronal communication. Additionally, the impact of polyplexes on axonal outgrowth in embryonic cortical neurons highlighted remarkable enhancements with Tg polyplexes carrying siPTEN compared to nTg polyplexes, showcasing their potential for promoting axonal development.

In summary, our developed Tg TMCSH-based polyplexes not only show promise for clinical translation but also hold therapeutic potential in gene silencing and promoting axonal growth. These findings collectively contribute to advancing the understanding of neural growth and regenerative therapies, opening avenues for innovative neurobiological applications. The Tg polyplexes, with their multifaceted attributes, present a robust platform for future investigations and developments in the field of neuron-Tg therapies.

## 4. Experimental methods

### 4.1. Trimethyl chitosan purification and preparation of thiolated trimethyl chitosan (TMCSH)

Trimethyl chitosan (TMC) was obtained from ultra-pure chitosan originating from *Agaricus bisporus* mushrooms (Kytozyme, lot VIHA0013-157, Belgium). The purification process for TMC involved filtration and dialysis, following a previously documented method [11]. Partially thiolated TMC (TMCSH) was obtained as previously described by us [16, 19]. The resulting TMCSH was stored at −20 °C until further usage.

### 4.2. Production, purification, and modification of the HC fragment

The HC fragment plasmid, supplied by Prof. Neil Fairweather from King’s College, UK, was produced using the BL21 *E. coli* strain and was subsequently purified following methods detailed in our previous study [20, 32]. The resulting purified protein fragment was modified with a bifunctional 5 kDa polyethylene glycol (PEG) spacer (Jenkem Technology, China) bearing an N-hydroxysuccinimide (NHS) and a maleimide (MAL) end group, at a 2.5 PEG per HC protein molar ratio [16]. The amount of reactive MAL groups in the HC fragment was determined through Ellman’s assay [44] and found to range between 1.0 and 1.2 mol PEG per HC.

### 4.3. Preparation of chitosan-based siRNA polyplexes

The preparation of polyplexes involved the use of siGFP or siPTEN [42, 45]. In cases where intracellular path monitoring was intended, a mimetic of siGFP (siRNAmi) labelled at the 5′ end of the sense strand with cyanine 5 (Cy5-siRNAmi) was used for polyplexes preparation. Integrated DNA Technologies supplied all sequences (see Table S1, SI). Briefly, TMCSH was dissolved overnight under stirring at room temperature (RT) in 5 mM hydrochloric acid (Fisher Scientific). Then, phosphate-buffered saline (PBS) 1× pH 7.4 (Sigma-Aldrich) was added, leading to a final pH 7.4 solution, which was kept under agitation for an additional 4 hours to complete polymer dissolution (TMCSH final concentration of 3 or 1 mg/mL). The resulting solution was filtered with a 0.22 μm pore-sized syringe filter (PES filter, Fisherbrand). Finally, the solution TMCSH final concentration was assessed using Cibacron® Brilliant Red (Sigma-Aldrich) as previously described by us [11]. The TMCSH-siRNA NP was allowed to stabilize for 15 minutes at RT before functionalization with the targeting molecule. Polyplexes were prepared following an N/P molar ratio (N/P 2, 4 and 6) [17–19]. Polyplexes between TMCSH and siRNA were prepared by combining siRNA (20 µM) with TMCSH solution in PBS 1×, pH 7.4 (siRNA final concentration 0.6 µM). To generate neuron-Tg NPs, HC-PEG (HC-PEG-MAL) was grafted into the NPs at a 4:1 HC-PEG to siRNA ratio (w/w) and incubated overnight with agitation (800 rpm) at RT [19].

### 4.4. Physicochemical characterization of non-targeted (nTg) and targeted (Tg) polyplexes

The nTg or Tg siGFP- or siPTEN-TMCSH polyplexes (N/P 2, 4 and 6) were prepared in PBS 1×, pH 7.4 and then physicochemically characterized. Namely we determined TMCSH complexation efficiency of siRNA (SYBR™ Gold assay and PAGE), polyplexes’ size and polydispersity index (DLS), surface charge (electrophoretic scattering), NP concentration in solution (NTA), and morphology (TEM).

#### 4.4.1. siRNA complexation efficiency

*SYBR™ Gold intercalation assay*. The siGFP or siPTEN complexation capacity was determined using the SYBR™ Gold assay, as previously outlined [42, 45]. The results are expressed as the relative percentage of complexation, with 100% denoting complete siRNA complexation, and 0% indicating the absence of any nucleic acid binding. Free TMCSH was employed to eliminate potential background fluorescence. *Polyacrylamide gel electrophoresis (PAGE)*. The ability of TMCSH to form complexes with siRNA was evaluated through gel electrophoresis. Polyacrylamide gels, composed of 4% (w/v) stacking and 15% (w/v) resolving gel, were prepared in Tris(tris(hydroxymethyl)aminomethane)/borate/EDTA (ethylenediaminetetraacetic acid) (TBE) 1× buffer (NZYtech). Polyplexes were prepared with TMCSH and siGFP or siPTEN at N/P ratios 2, 4, and 6, as detailed above. An equivalent to 6 pmol siPTEN was loaded into each well. After electrophoresis (run at 100 V for 30 minutes), the gel was treated with 1× SYBR™ Gold (Invitrogen) nucleic acid staining solution for 10 minutes and visualized using a GelDoc XR imager (Bio-Rad Laboratories). The intensity of the bands was quantified using Image Lab software (version 6.1.0, Bio-Rad Laboratories). The band intensity was compared to that of the control (free form siPTEN) and expressed as the relative percentage of free siRNA, where 100% indicates that the nucleic acid is completely unbound.

#### 4.4.2. Hydrodynamic size and zeta potential measurements

The hydrodynamic size and PDI of the formed Tg and nTg polyplexes were assessed following established protocols using a Malvern Zetasizer Nano ZS (Malvern Instruments, UK) [42]. Analysis was conducted using the Zetasizer Software (version 7.13).

#### 4.4.3. Polyplexes’ stability studies

Tg N/P 4 polyplexes’ stability was explored in the different cell culture media explored in this study through DLS. Polyplexes solutions were six times diluted in the different cell culture media and incubated for 4 hours at 37 °C environment with 5% CO_2_. Polyplexes solution diluted in PBS 1× was used as controls. All samples were at pH 7.4.

#### 4.4.4. Nanoparticle tracking analysis (NTA)

Tg siPTEN-polyplexes at an N/P ratio 4 were prepared following the procedure previously outlined. These polyplexes were then diluted in PBS 1× to a final volume of 1 mL (concentration range of 10^7^–10^9^ particles per mL). Samples were tracked and analysed in a NanoSight NS300 equipped with a 488 nm blue laser using previously established methods [42]. The raw data obtained from laser scattering and particle movement underwent analysis via NTA software (version 3.4; NanoSight Ltd, UK). Automatic settings were applied, including minimum expected particle size, blur, and minimum track length. The detection threshold was fixed at 4, while the sample viscosity was adjusted to match the water’s viscosity (at 25 °C), and the temperature was set to 25 °C. The output data was represented as particle concentration, measured in the number of particles per mL.

#### 4.4.5. Transmission electron microscopy (TEM)

Tg siRNA-polyplexes were prepared at N/P 4 and imaged following our established protocol using a Jeol JEM 1400 operating at 80 kV [45]. Subsequently, ImageJ software (version 1.54d; National Institutes of Health, USA) was used to process the acquired images.

### 4.5. Animals and cell culture

The culture of cell lines, including ND7/23 (mouse neuroblastoma and rat neuron hybrid, ECACC), HT22 (mouse hippocampal neuronal, provided by Dave Schubert at the Salk Institute), and NIH 3T3 (mouse embryonic fibroblast, ECACC), was conducted using Dulbecco’s Modified Eagle’s Medium (DMEM) supplemented with GlutaMAX™ (Gibco). The culture medium consisted of 10% (v/v) heat-inactivated foetal bovine serum (FBS) (56 °C for 30 minutes) and 1% (v/v) antibiotic solution containing 10,000 U/mL penicillin and 10,000 μg/mL streptomycin (P/S), all from Gibco. These cells were maintained in a 37 °C environment with 5% CO_2_. Regular polymerase chain reaction (PCR) analyses were performed to ensure the absence of mycoplasma contamination. The cells used for the experiments were between passages 1 and 15 post-thawing from liquid nitrogen. Animal experiments followed European Union guidelines (EU Directive 2010/63/EU) and Portuguese law (DL 113/2013) with a priority on minimizing animal suffering and the total number of animals used. All procedures involving animals were approved by the official Portuguese authority on animal welfare and experimentation (*Direção Geral de Alimentação e Veterinária* - DGAV), Institute for Research and Innovation in Health (i3S) Ethics Committee (CEA, i3S), and the Animal Ethics Committee of Tel Aviv University (TAU). The animals were housed in enriched environments with unrestricted access to food and water, maintaining a 12/12-hour light/dark cycle, and controlled ambient temperature and humidity at i3S or TAU facilities. DRG, motor, and cortical neuron cultures were obtained from mouse embryos (E16.5, E12.5 and E16.5, respectively) originating from pregnant 2–3-month-old female C57BL/6 mice. For downregulation studies, HB9::GFP mice obtained from Jackson Laboratories were used and maintained by breeding with ICR mice. Euthanasia procedures for mice followed ethical guidelines and institutional protocols. Initially, mice were placed in a CO_2_ chamber until unconsciousness, which was followed by cervical dislocation. Embryonic DRG explants were collected from E16.5 wild-type C57BL/6 embryos. The embryos were carefully harvested and kept in ice-cooled Hank’s balanced salt solution (HBSS, pH 7.4, Sigma) without CaCl_2_ and MgSO_4_ until the DRGs were extracted. Once the spinal cord was removed, access to the embryonic ganglia was achieved via the dorsal side. DRGs were isolated under a stereomicroscope and seeded on coverslips (µ-Slide 15 Well 3D Glass Bottom, #81507, ibidi, Martinsried, Germany). The DRG explants were cultured in Neurobasal medium (Gibco), supplemented with 2% (v/v) B-27 (Thermo Fisher Scientific), 2 mM L-glutamine (Gibco), and 25 ng/mL nerve growth factor (NGF, Millipore).

The motor neuron cultures were isolated from E12.5 HB9::GFP C57BL/6 embryos following the procedure described by Ionescu et al. (2022) [46]. Cells were cultured in Neurobasal medium (Gibco) supplemented with B27 1× (Gibco), 2% (v/v) horse serum (Gibco), 1% (v/v) P/S, 0.025 mM GlutaMAX™, 25 µM beta-mercaptoethanol (Sigma), 1 µM glial cell line-derived neurotrophic factor (GDNF), 0.5 µM ciliary neurotrophic factor (CNTF), and 1 µM brain-derived neurotrophic factor (BDNF) (all factors from Alomone Labs). Primary mouse neuronal cortical cells were isolated from E16.5 wild-type C57BL/6 embryos’ prefrontal cortex. Cerebral cortices were dissected, then incubated with trypsin (1.5 mg/mL, Gibco) in HBSS without CaCl_2_ and MgSO_4_ at 37 °C for 12 minutes. Following incubation, 10% (v/v) FBS-containing HBSS was added. The resultant cell clusters underwent gentle washing three times with non-supplemented HBSS, followed by the addition of the completed Neurobasal medium (previously described). Following mechanical dissociation, the cell suspension was filtered using a 70 µm nylon cell strainer (Corning Falcon™), then seeded (cell density specified in the respective subsection) and incubated at 37 °C in 5% CO_2_. The purity of the cortical neuron culture was assessed through immunocytochemistry (data not shown) using specific antibodies: rabbit anti-glial fibrillary acidic protein (GFAP), rat myelin basic protein (MBP), rabbit anti-oligodendrocyte transcription factor 2 (Olig2), and rabbit anti-ionized calcium-binding adapter molecule 1 (IBA1). The culture’s purity, determined through image analysis using Ilastik and Cell Profiler software, was found to be over 97%.

### 4.6. Biocompatibility evaluation in well-plates

The effect of Tg and nTg TMCSH-based polyplexes on embryonic mice motor neurons and cortical neurons was examined using LDH assay and MEAs. A day before cell seeding, the well-plates were coated with poly(L-ornithine) (PLO, 1.5 µg/mL in PBS 1× pH 7.4, Sigma-Aldrich) and laminin (5 μg/mL in autoclaved water, Sigma) for primary motor neurons, or poly(D-lysine) (PDL, 50 µg/mL in 0.1 M borate buffer pH 8.5, Sigma-Aldrich) for primary cortical neurons.

#### 4.6.1. LDH assay

After coating, 20 × 10^4^ primary motor or cortical viable neurons (trypan blue assay) were seeded into each well. The cells were cultured in an incubator set at 37 °C and 5% CO_2_, with 50% of the medium refreshed every 2 days. At day *in vitro* (DIV) 5, both Tg and nTg polyplexes, at N/P 4, and complexing siPTEN, were introduced at a final siPTEN concentration of 100 nM. Cells were incubated for 24 hours, the cell culture medium collected, and the released LDH quantified using the CyQUANT™ LDH cytotoxicity assay kit (Thermo Fisher), as previously described by us [42]. As part of the controls, cells were treated with Lipofectamine 2000 (L2k)-siPTEN lipoplexes, and Triton X-100 0.1% (v/v, Invitrogen) in PBS 1× pH 7.4. The data is presented as the percentage of LDH released by treated cells concerning cells not exposed to the treatment (negative control) [42].

#### 4.6.2. Neuronal electrical activity

To monitor the electrical activity of neuronal networks, commercial six-well-plate MEAs (60-6wellMEA200/30iR-Ti, Multichannel Systems) were used. Each well contained a recording area with a 3 × 3 grid of embedded gold electrodes (30 µm diameter, spaced 200 µm apart), allowing non-invasive monitoring. A day before cell seeding, the well-plates were coated as mentioned in the previous section. Subsequently, 20 × 10^4^ primary motor or cortical viable neurons (trypan blue assay) were seeded into each well. The cells were cultured in an incubator set at 37 °C and 5% CO_2_, with 50% of the medium refreshed every 2 days. At DIV5, N/P 4 siPTEN-polyplexes were introduced to the cells (final siRNA concentration of 100 nM). At DIV14, spontaneous neuronal activity was recorded using the MEA setup (sampling rate of 10 kHz, recording duration of 5 minutes). Cells were treated with Triton X-100 0.1% (v/v) in PBS 1× pH 7.4 were used as control. The detected spikes were sorted and analysed using a MATLAB script (R2024a, The MathWorks, Inc., Natick, MA, USA), Github (https://github.com/MaozLab/MaozAnalyzer).

### 4.7. Neurospecificity, internalization and transfection evaluation

#### 4.7.1. Cellular binding and neurospecificity in cell lines

The Tg polyplexes’ interaction with ND7/23, HT22, and NIH 3T3 cells was evaluated using flow cytometry. Initially, cells were seeded in 24-well plates at a density of 2.5 × 10^4^ viable cells (trypan blue assay) per cm^2^, as described above. Tg polyplexes, at N/P 2, 4, and 6 and complexing Cy5-siRNAmi, were introduced at a final siRNAmi concentration of 100 nM. Following 8 hours of incubation, the cells underwent trypsinization, PBS 1× rinsing, centrifugation, and resuspension in PBS 1× with 2% (v/v) FBS. The subsequent analysis was performed using a BD Accuri™ C6 flow cytometer (BD Biosciences). The Cy5 fluorescence signal was measured to quantify the positive cells (10,000 viable cells per sample). Untreated cells and cells treated with Lipofectamine RNAiMAX (L-iMax, Invitrogen) carrying Cy5-siRNAmi were used as controls. Data analysis was conducted using FlowJo software (version 10, FLOWJO, LLC).

#### 4.7.2. Membrane receptor blocking assay

ND7/23, HT22, and NIH 3T3 cells were seeded in 24-well plates at a density of 3.75 × 10^4^ viable cells (trypan blue assay) per cm^2^. Twenty-four hours after seeding, cells were incubated at 4 °C for 15 minutes, followed by three rinses with cold HBSS without Ca^2+^ and Mg^2+^. Subsequently, cells were incubated with the free form of HC (HC final concentration 1 nM) in HBSS with 0.1% (w/v) bovine serum albumin (BSA, NZYTech) for 15 minutes at 4 °C. After HC incubation, cells were treated with Tg and nTg N/P 4 polyplexes for 8 hours at 37 °C, 5% CO_2_ and finally analysed via flow cytometry (as detailed above).

#### 4.7.3. Internalization capacity explored in 3D ex vivo explants

After removal, embryonic DRG explant cultures remained undisturbed for 72 hours. Tg and nTg N/P 4 polyplexes carrying Cy5-siRNAmi were prepared as previously detailed. Transfection was performed using an 8 μL polyplex suspension within a final volume of 50 μL of culture medium (nucleic acid final concentration of 100 nM), and explants were incubated in the presence of NPs for 24 hours. Subsequently, explants were washed with PBS 1× and fixed with paraformaldehyde (PFA). The fixation process was performed in three stages: first, PFA was added to the cell medium in the well at a 1:1 ratio and incubated for 15 minutes. Next, the solution was removed, and 50 μL of fresh PFA was added and incubated for another 15 minutes. After incubation, the explants were carefully washed three time with PBS 1×. Following fixation, explants were treated with a permeabilization buffer (Triton™ X-100, 0.3% (v/v) in PBS 1×) for 45 minutes, followed by incubation in a blocking buffer composed of 1% (w/v) BSA in PBS 1× for 45 minutes at RT. After carefully rinsing explants with PBS 1×, they were incubated overnight at 4 °C with mouse anti-βIII tubulin (2 µg/mL, TUBB3, Biolegend). After the primary antibody incubation, samples were treated with Alexa Fluor 488 donkey anti-mouse IgG (H+L) secondary antibody (2 µg/mL, A-21202, Invitrogen) for 1 hour at RT and then washed again with PBS 1×. Cell nuclei were then stained using a Hoechst 33 342 solution (5 µg/mL, Thermo Fisher Scientific) in PBS 1× for 10 minutes. Samples were left in PBS 1× at 4 °C until further use. Control experiments where DRGs were left untreated or were treated with free nucleic acid (Cy5-siRNAmi) or a TMCSH solution were conducted. The Leica SP8 Confocal Microscope (Leica Microsystems, Inc), equipped with an HC PL APO CS2 10x/0.40 dry objective, was used to capture images of the samples. Three-dimensional z-stacks were acquired at a resolution of 2048 × 2048 pixels (zoom 1; pixel size: 0.568 µm in the x and y axes, and 5 µm in the z-axis). After the acquisition, image processing was carried out using the Leica Application Suite X software (version 3.5.7.23225, Leica Microsystems CMS GmbH).

#### 4.7.4. Downregulation of reporter gene expression in neurons

Primary mouse motor neurons were obtained from E12.5 HB9::GFP mouse embryo ventral spinal cord horns [31]. HB9::GFP motor neurons were seeded onto 24-well plates (coated with 150 µg/mL PLO and 3 µg/mL laminin) at a density of 2.5 × 10^4^ viable cells (trypan blue assay) per cm^2^. These cells were incubated in supplemented medium. Polyplexes containing siGFP at N/P 4 were prepared as detailed above. Transfection was carried out using 50 μL polyplexes in 300 μL of culture medium (final siGFP concentration of 100 nM). Untreated cells and cells treated with lipoplexes carrying siPTEN were used as controls. Twenty-four hours post-treatment, the medium was replaced by fresh supplemented medium, and cells were further incubated for 72 hours. Subsequently, the cells were rinsed with PBS 1× and fixed for 15 minutes using PFA (4% (w/v) in PBS 1×). Following cell fixation, cells were treated with a permeabilization buffer (Triton™ X-100, 0.1% (v/v) in PBS 1×) for 30 minutes. Cell nuclei were stained using Hoechst 33 342 in PBS 1× for 10 minutes. Coverslips were mounted using VectaShield (Vector Laboratories), and image acquisition was conducted using an Olympus FV3000 confocal microscope equipped with 20×/0.8, and 60×/1.42 NA objectives. Confocal images of 512 × 512 pixels (zoom 1, 0.622 µm per pixel) were analysed using ImageJ (version 1.54d).

### 4.8. Intracellular trafficking and dynamics

The selected microfluidic chips had two compartments interconnected through a network of 139 microchannels (2 × 10 μm) with a length of 450 µm, allowing axons to extend within them. The chip was produced using a prepolymer mixture of SYLGARD™ 184 Silicone Elastomer (10:1 mix of silicone elastomer and its curing agent, Dow Corning). It was cast and cured for 2 hours at 70 °C, against the positive master mold to generate a negative replica mold. Following the curing process, poly(dimethylsiloxane) (PDMS) devices were separated from the master mold, and the media reservoirs were created using a steel biopsy punch (Ø 5 mm, KAI medical, BP-50F). After sterilization, achieved by washing with 70% (v/v) ethanol and subjecting the devices to 10 minutes of exposure to ultraviolet light, PDMS platforms were plasma-treated using air (Tergeo plasma cleaner TG100, Pie Scientific) and affixed onto coated glass coverslips (22 × 22 mm, Normax), lightly pressing both surfaces. The glass coverslips were previously coated as described above.

Motor or cortical neurons were seeded into the proximal compartment (cell soma compartment) of the two-compartment microfluidic device (Figure 3A) at a density of 25 × 10^4^ viable cells (trypan blue assay) in 0.161 cm^2^ (area of compartment and reservoirs). To create a hydrostatic pressure difference and enable a slow but continuous flow through the microchannels, an approximate volume difference of 50 μL was maintained between the proximal and distal compartments (representing the cell soma and axonal compartments, respectively), with the lower volume allocated to the axonal compartment. This setup facilitates the fluidic isolation of the cell soma and axonal compartments.

At DIV5, the flow was applied to the culture using an Ismatec IPC peristaltic pump (Cole-Parmer, USA, flow rate: 56.5 µL/min). Tg and nTg N/P 4 polyplexes, carrying Cy5-siRNAmi, were added into the reservoir with the medium circulating through the axonal compartment. Upon addition to the cultures, polyplexes only contacted with the axonal terminals (refer to Figure 3A, compartment 2) in order to evaluate their ability to interact, get internalized, and reach the neuronal cell body. Untreated cells were used as the control group. Twenty-four hours after introducing the polyplexes into the system under flow, it was disassembled, and the cells were fixed with PFA in a microtubule-protecting (MP) fixative buffer. MP-PFA solution consists of 4% (w/v) PFA, 65 mM 1,4-Piperazinediethanesulfonic acid (PIPES), 25 mM 4-(2-hydroxyethyl)-1-piperazineethanesulfonic acid (HEPES), 10 mM ethylene glycol-bis(β-aminoethyl ether)-N,N,N′,N′-tetra acetic acid (EGTA), and 3 mM MgCl_2_ in PBS 1× (all acquired from Sigma-Aldrich). The fixation process was conducted in three stages: initially, MP-PFA was added to the cell medium in the microfluidic reservoirs (1:1 ratio) and left to incubate for 15 minutes. Following this, the solution was removed, and fresh MP-PFA was added to fully cover the reservoirs and incubated for 15 minutes. Finally, the MP-PFA from the reservoirs was removed, the microfluidic device was separated from the coverslip, and MP-PFA was directly applied to the coverslip, which was incubated for an additional 15-minute period. Following cell fixation, the cells were treated with a permeabilization buffer (Triton™ X-100, 0.1% (v/v) in PBS 1×) for 15 minutes, followed by incubation in a blocking buffer composed of 1% (w/v) BSA (NZYTech) in PBS 1× for 45 minutes at RT.

After rinsing the cells with PBS 1×, they were incubated overnight at 4 °C with mouse anti-βIII tubulin (2 µg/mL). After the primary antibody incubation, samples were treated with Alexa Fluor 488 donkey anti-mouse IgG (H+L) secondary antibody (2 µg/mL) for 1 hour at RT and then washed again with PBS 1×. To conclude, cell nuclei were stained using a Hoechst 33 342 solution (5 µg/mL, Thermo Fisher Scientific) in PBS 1× for 10 minutes. Fluoromount™ Aqueous Mounting Medium (Sigma) was applied to mount the coverslips on the microscope slides. Image capture was performed using a Leica TCS SP5 Confocal Microscope (Leica Microsystems, Inc.) equipped with an HCX PL APO CS 63.0x1.40 oil objective. Confocal image z-stacks of 1024 × 1024 pixels (16-bit depth and zoom levels between 1.7 and 2.5, voxel size: 240.5 nm) were obtained. For Hoechst, Alexa Fluor 488, and Cy5 acquisition, the respective SP mirror channels were set to 414-486 nm, 497-589 nm, and 695 -765 nm. Subsequently, the images were analyzed using Leica Application Suite X software (Leica Microsystems) and Fiji ImageJ (version 1.54f).

#### 4.8.1. Intracellular dynamics assessment

To assess the retrograde axonal transport of the TMCSH-siRNA NPs, motor or cortical neurons were initially seeded in the proximal compartment (cell soma compartment) of a two-compartment microfluidic setup (Figure 3A) at a density of 25 × 10^4^ viable cells (trypan blue assay) in 0.161 cm^2^. The seeding procedure was previously detailed. At DIV5, Tg and nTg N/P 4 polyplexes carrying Cy5-siRNAmi were added into the axonal compartment (final siRNAmi concentration of 100 nM). Following a 16-hour incubation period, live axonal imaging was performed using a Crest X-Light V3 spinning disk confocal microscope (Nikon) equipped with a 37 °C heated chamber. At least 24 microchannels, each containing a minimum of one axon, were randomly selected for live cell imaging within each treatment group. Thirty minutes before image acquisition, cells were incubated with CellMask™ Plasma Membrane Stain 488 (1:1000 dilution, C37608, Invitrogen). Images were captured for a minimum duration of 5 minutes at a frame rate of 0.33 frames-per-second (0.33 FPS) with a resolution of 1024 × 1024 pixels, using a Plan Apo VC 60x A WI DIC N2 water objective (numerical aperture: 1.2). Data were collected from three independent experiments, and subsequent image analysis was performed using Fiji ImageJ 1.54f as described by Ionescu et al. (2022) [46].

To delve deeper into intracellular dynamics, the process of cell seeding and treatment with polyplexes was done as mentioned above, but the incubation time with the NPs was extended. At specific intervals (8, 12, 24, 48, and 120 hours), cells were fixed with MP-PFA using the three-step fixation method, immunostained and imaged as formerly detailed.

### 4.9. Biocompatibility profile of polyplexes assessed in microfluidic neuronal cultures

#### 4.9.1. LDH assay

Upon seeding cells, the cells were subjected to nTg or Tg N/P 4 polyplexes, prepared with siPTEN. Polyplexes were added in the axonal compartment. Following a 24-hour exposure, the cell culture medium from the 5 mm cell soma compartments was collected, and the released LDH was quantified as outlined earlier.

#### 4.9.2. Neuronal electrical activity evaluation using MEAs-integrated microfluidic platforms

Commercial planar MEAs (60MEA200/30iR-ITO, Multichannel Systems) featuring sixty electrodes (30 µm in diameter, spaced 200 µm apart) were used to monitor the electrical activity of neuronal networks within two-compartment microfluidic setups (Figure 3A). Before use, both the microfluidic devices and MEAs underwent a brief submersion in 70% ethanol, followed by air-drying in a laminar flow hood for sterilization. Subsequently, the microfluidic devices were meticulously assembled on top of the MEAs under stereomicroscope guidance to ensure precise alignment of the microchannels with the microelectrodes. The platform was then coated according to the procedure previously described. The procedure involved seeding primary motor neurons or cortical neurons (20 × 10^4^ viable cells) as described in this section. Throughout the experiment, a maintained hydrostatic pressure difference ensured appropriate axonal growth direction. At DIV5, N/P 4 polyplexes were added to cells in the axonal compartment (Figure 3A, compartment 2). On DIV14, spontaneous neuronal activity in the cell body compartment was recorded using the same settings. Microfluidic systems with untreated cells and cells treated with Triton 0.1% (v/v) were control groups. Detected spikes were sorted and analysed using a MATLAB script and SpikeSorter software (version 5.20, Vancouver, BC Canada) [47].

### 4.10. Axonal outgrowth

To evaluate the biological impact by studying axonal outgrowth, a three-compartment microfluidic system with two sets of 600 µm-length microchannels was employed (Figure 3B). Primary cortical neurons were seeded in the central compartment of this three-compartment microfluidic setup (Figure 3B, compartment 2) at a density of 20 × 10^4^ viable cells (trypan blue assay) in 0.076 cm^2^ (area of central compartment and reservoirs). On DIV5, Tg and nTg N/P 4 polyplexes containing siPTEN were added in the cell soma compartment (Figure 3B, compartment 2, final siRNA concentration of 100 nM). Cells treated with free TMCSH, siPTEN, and untreated cells were used as controls. At DIV10, the cells were fixed using MP-PFA, following the 3-step fixation procedure described above. After cell fixation, cells were treated with permeabilization buffer (Triton™ X-100, 0.1% (v/v) in PBS 1×) for 15 minutes, followed by an hour-long incubation in a blocking buffer composed of 1% BSA in PBS 1× at RT. The cells were then rinsed with PBS 1× and incubated overnight at 4 °C with rabbit anti-Neurofilament H antibody (0.6 µg/mL, ab207176, Abcam). After the primary antibody incubation, samples were treated with Alexa Fluor 488 donkey anti-rabbit IgG (H+L) secondary antibody (2 µg/mL, A-21206, Invitrogen) for an hour at RT and washed again with PBS 1×. Subsequently, cell nuclei were stained using Hoechst 33 342 (5 µg/mL, Thermo Fisher Scientific) in PBS 1× for 10 minutes. Finally, Fluoromount™ Aqueous Mounting Medium (Sigma) was applied to mount the coverslips. The entire microfluidics was imaged in a Leica DMI6000 using HCX PL FLUOTAR L 20x/0.40 CORR Ph1 and HCX PL FL L 40x/0.60 CORR Ph2 objectives. Images of 512 × 512 pixels (zoom 1, voxel size of 1 µm). The axonal outgrowth within the axonal compartments (Figure 3B, compartments 1 and 3) was assessed using linear Sholl analysis MATLAB scripts, kindly provided by Martin Offterdinger and Prof. Rüdiger Schweigreiter [43].

### 4.11. Statistical data analysis

Analysis and presentation of data were conducted using IBM SPSS Statistics version 26 (SPSS Inc., USA) and GraphPad Prism version 9 for Windows (GraphPad Software, USA). The normal distribution of data was verified using the Shapiro-Wilk test. Differences were assessed through parametric one-way or two-way ANOVA, followed by Tukey’s multiple comparisons tests, as detailed in the results. Statistical significance was denoted by P values less than 0.05.

## Conflicts of interest

The authors declare that there are no potential conflicts of interest that could have influenced the work presented in this paper.

## Data availability statement

The data supporting the findings of this study can be obtained from the corresponding author upon reasonable request.

## Supporting information

Supporting Information is available from the Wiley Online Library or from the author.

## Supporting information

Support Information

## Acknowledgements

This work was supported by Portuguese funds through *Fundação para a Ciência e a Tecnologia*, I. P. (FCT) in the framework of the projects PTDC/CTM-NAN/3547/2014 and PTDC/BTM-MAT/4156/2021. A. P. S. acknowledges FCT for her Ph.D. scholarship (SFRH/BD/137073/2018), EMBO for her Scientific Exchange Grant (10398) and COST for her short-term scientific mission grant (9f51e2a4, COST Action: CA17103 -Delivery ofAntisense RNA ThERapeutics (DARTER) Action). V. L. appreciates her Assistant Researcher contract under the “Concurso Estímulo ao Emprego Científico Individual – 4ª Edição” (2021.00472.CEECIND), and S. C. G. her post-doctoral fellowship (SFRH/BPD/122920/2016). B. M. M. also acknowledges the support of the Israel Science Foundation (ISF grant: 2248/19, 1934/23), ERC SweetBrain 851765, The Aufzien Family Center for the Prevention and Treatment of Parkinson’s Disease at Tel Aviv University, The Zimin Foundation, the Innovation Authority “BioChip” and the Israel Ministry of Science and Technology (Grant No. 3–17351, 8-2175), the European Union Horizon 2020 research and innovation programme under the Marie Skłodowska-Curie (grant agreement No. 101007804-Micro4Nano) and a OSTEONET project grant (agreement No 101086329). The size and zeta potential analyses were conducted at the Biointerfaces and Nanotechnology Scientific Platform, with the assistance of Ricardo Vidal. Confocal microscopy was conducted at the i3S Scientific Platforms Bioimaging, with the assistance of Dr María Lázaro, and at Advanced Light Microscopy, with the assistance of Dr Maria Azevedo and Dr Paula Sampaio. These platforms are members of the Portuguese Platform of Bioimaging (PPBI-POCI-01-0145-FEDER-022122).

## Notes

### Competing Interest Statement

The authors have declared no competing interest.

