## Supplementary material for "Engineered chitosan-derived nanocarrier for efficient siRNA delivery to peripheral and central neurons": Support Information

A. P. Spencer
i3S – Instituto de Investigação e Inovação em Saúde, Universidade do Porto, Porto, Portugal
INEB – Instituto de Engenharia Biomédica, Universidade do Porto, Porto, Portugal
Faculdade de Engenharia, Universidade do Porto, Porto, Portugal

A. Vilaça, M. Xavier
International Iberian Nanotechnology Laboratory (INL), Braga, Portugal

R. Santos, M. Lázaro, V. Leiro, S. C. Guimarães
i3S – Instituto de Investigação e Inovação em Saúde, Universidade do Porto, Porto, Portugal
INEB – Instituto de Engenharia Biomédica, Universidade do Porto, Porto, Portugal

A. Ionescu, E. Perlson
Sagol School of Neuroscience, Tel Aviv University, Tel Aviv, Israel
Department of Physiology and Pharmacology, Sackler Faculty of Medicine, Tel Aviv University, Tel Aviv, Israel

B. M. Maoz
Sagol School of Neuroscience, Tel Aviv University, Tel Aviv, Israel
Department of Biomedical Engineering, Tel Aviv University, Tel Aviv, Israel
The Center for Nanoscience and Nanotechnology, Tel Aviv University, Tel Aviv, Israel
Sagol Center for Regenerative Medicine, Tel Aviv University, Tel Aviv, Israel

A. P. Pêgo
i3S – Instituto de Investigação e Inovação em Saúde, Universidade do Porto, Porto, Portugal
INEB – Instituto de Engenharia Biomédica, Universidade do Porto, Porto, Portugal
Instituto de Ciências Biomédicas Abel Salazar (ICBAS), Universidade do Porto, Porto, Portugal

* Corresponding authors

### Table of Contents

Page


| | | |
|---|---|---|
| 1) | siRNA sequences | 3 |
| 2) | Physicochemical features of TMCSH-based polyplexes | 4 |
| 3) | Stability studies of neuron-targeted polyplexes | 5 |
| 4) | Biocompatibility evaluation in well-plates | 6 |
| 5) | Cellular association of non-targeted TMCSH-derives nanoparticles | 9 |
| 6) | Axonal transport of neuron-targeted polyplexes – Kymographic analysis | 10 |
| 7) | Biocompatibility profile of polyplexes assessed in microfluidic neuronal cultures | 11 |

#### 1) siRNA sequences

**Table S1.** Nucleic acids used in this study and their correspondent sequences.

| Nucleic acid | Sequence (5' – 3') |
|---|---|
| siPTEN | sense: CGACUUAGACUUGACCUAUAU<br>antisense: AUAGGUCAAGUCUAAGUCGAA |
| siGFP | sense: GCUGACCCUGAAGUUCAUCUGCACC<br>antisense: GGUGCAGAUGAACUUCAGGGUCAGCUU |
| Cy5-siRNAmi | sense: GGTGCAGATGAACTTCAGGGTCAGCTT<br>antisense: /Cy5/GCTGACCCTGAAGTTCATCTGCACC |

2'-*O*-methyl RNA bases are underlined

#### 2) Physicochemical features of TMCSH-based polyplexes

**Table S2.** Physicochemical characterization of non-targeted (nTg) and targeted (Tg) TMC-based nanoparticles carrying siGFP or siPTEN prepared at N/P ratios of 2, 4 and 6, in phosphate-buffered buffered saline (PBS) 1× pH 7.4. Complexation efficiency of siRNAs was determined by the SYBR™ Gold assay, average hydrodynamic size and polydispersity index (PDI) by dynamic light scattering (DLS), and zeta potential by electrophoresis at 25 °C. Values are expressed as mean ± SD (standard deviation) of three independent experiments (n=3), with one replicate per experiment. For statistical analysis, one-way ANOVA test was used. Significant differences: ****p ≤ 0.0001.

| siRNA | Formulation | N/P ratios | Complexation efficiency (%) | Hydrodynamic size (nm) | PDI | Zeta potential (mV) |
|---|---|---|---|---|---|---|
| siGFP | nTg | 2 | 63.7 ± 3.1 | 176.4 ± 3.0 | 0.01 ± 0.01*** | 13.8 ± 1.9 |
| | | 4 | 65.2 ± 3.1 | 134.3 ± 5.3** | 0.14 ± 0.03 | 14.2 ± 1.3 |
| | | 6 | 69.6 ± 1.5 | 169.0 ± 5.9 | 0.15 ± 0.03 | 13.1 ± 1.9 |
| | Tg | 2 | 75.3 ± 2.4 | 203.4 ± 25.4** | 0.06 ± 0.02*** | 12.0 ± 1.6 |
| | | 4 | 75.2 ± 1.7 | 155.9 ± 5.8 | 0.17 ± 0.01 | 13.8 ± 1.0 |
| | | 6 | 74.5 ± 3.6 | 165.2 ± 18.3 | 0.14 ± 0.02 | 13.2 ± 1.2 |
| siPTEN | nTg | 2 | 61.5 ± 0.3 | 197.4 ± 14.4* | 0.01 ± 0.01*** | 7.8 ± 1.3** |
| | | 4 | 66.3 ± 4.0 | 150.8 ± 8.6 | 0.14 ± 0.03 | 12.8 ± 1.5 |
| | | 6 | 72.4 ± 3.2 | 166.4 ± 10.1 | 0.22 ± 0.04 | 12.6 ± 1.5 |
| | Tg | 2 | 66.7 ± 4.2 | 177.7 ± 1.5 | 0.05 ± 0.02*** | 9.3 ± 1.2 |
| | | 4 | 68.2 ± 1.8 | 130.5 ± 8.2* | 0.17 ± 0.03 | 13.4 ± 1.9 |
| | | 6 | 74.1 ± 0.8 | 176.2 ± 6.22 | 0.25 ± 0.04 | 12.2 ± 1.0 |

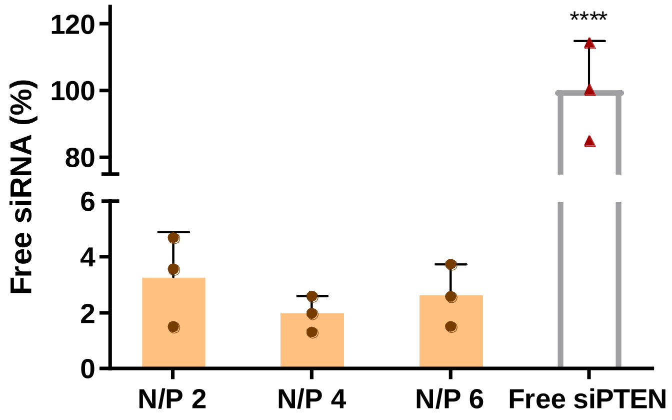


**Figure S1.** Intensity of bands in polyacrylamide gel electrophoresis (PAGE), compared to the intensity of free siRNA band. PAGE was done to assess the Tg polyplexes complexation capacity of siPTEN (Figure 1B). Polyplexes were prepared in PBS 1× pH 7.4. Results are shown as mean ± SD of three independent experiments (n = 3), with one replicate per experiment. For statistical analysis, one-way ANOVA test was used. Significant differences: ****p ≤ 0.0001.

#### 3) Stability studies of neuron-targeted polyplexes

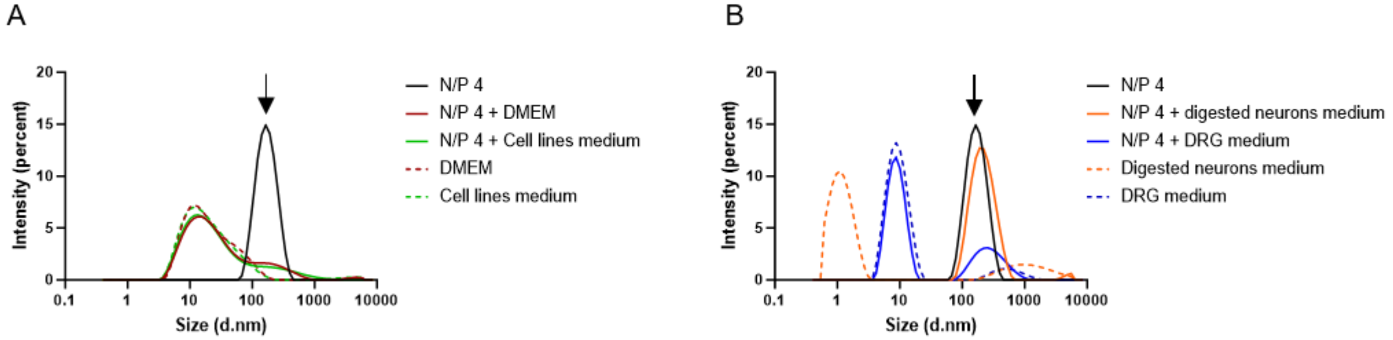


**Figure S2.** Representative size distributions (Dynamic Light Scattering, DLS) of targeted N/P 4 siPTEN-dendriplexes diluted in PBS 1x (dark blue line, indicated with a black arrow), in (A) Dulbecco's Modified Eagle's Medium (DMEM) supplemented with GlutaMAX™ (red line), DMEM supplemented with 10% (v/v) heat-inactivated fetal bovine serum (FBS) and 1% (v/v) penicillin and streptomycin (P/S) (green line), in (B) Neurobasal medium, supplemented with 2% (v/v) B-27, 2 mM L-glutamine, and 25 ng/mL nerve growth factor (NGF) (blue line) and in Neurobasal medium supplemented with B27 1×, 2% (v/v) horse serum, 1% (v/v) P/S, 0.025 mM GlutaMAX™, 25 µM beta-mercaptoethanol, 1 µM glial cell line-derived neurotrophic factor (GDNF), 0.5 µM ciliary neurotrophic factor (CNTF), and 1 µM brain-derived neurotrophic factor (BDNF) (orange line). All buffers and media were at pH 7.4.

#### 4) Biocompatibility evaluation in well-plates

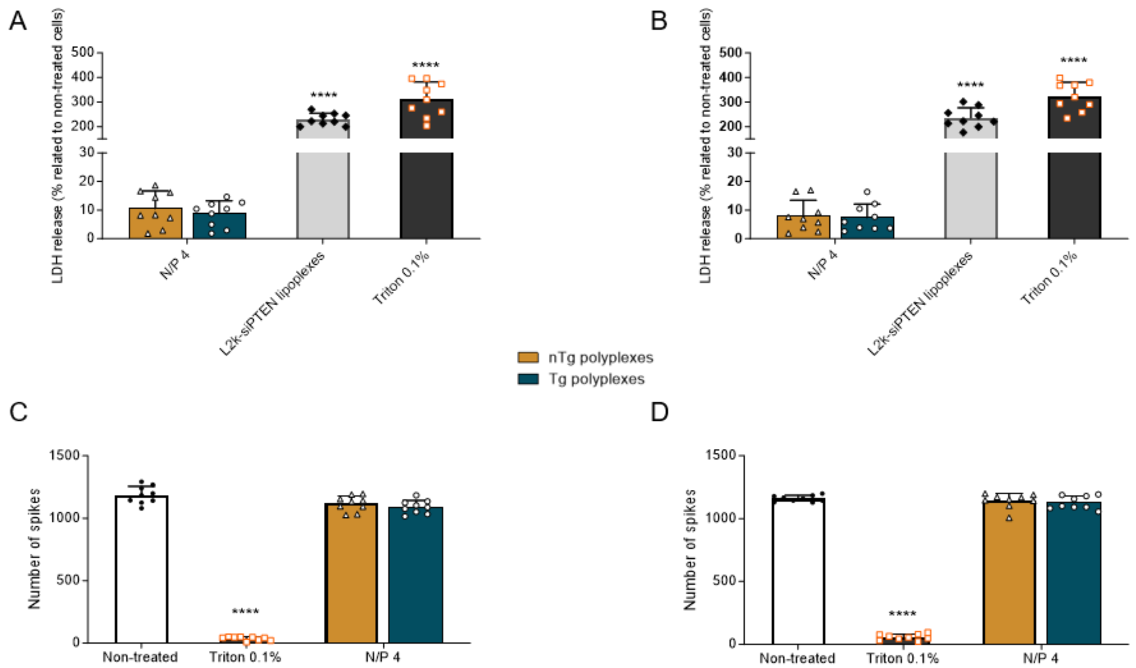


**Figure S3.** Biocompatible profile evaluation though the evaluation of plasma membrane integrity of A) motor neurons and B) cortical neurons seeded in well-plates, assessed by LDH assay after 24 hours of incubation with polyplexes. Results are shown as mean ± SD of three independent experiments (n = 3) with three replicates per experiment. For statistical analysis, two-way ANOVA test was used. Significant differences: ****$p \leq 0.0001$, *versus* non-treated cells.

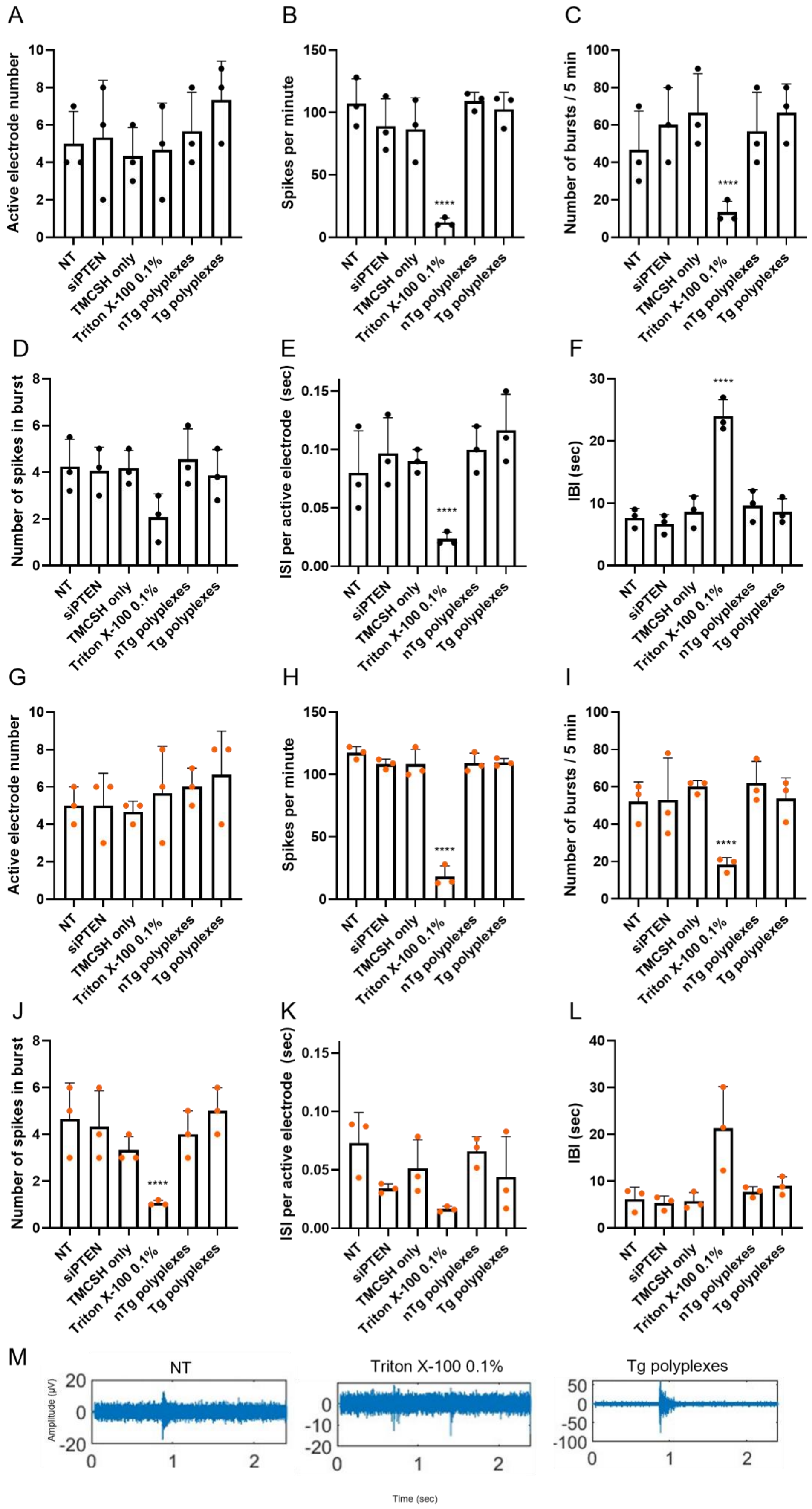
A
B
C
D
E
F
G
H
I
J
K
L
M
Active electrode number
Spikes per minute
Number of bursts / 5 min
Number of spikes in burst
ISI per active electrode (sec)
IBI (sec)
NT
siPTEN
TMCSH only
Triton X-100 0.1%
nTg polyplexes
Tg polyplexes
****
Triton X-100 0.1%
Tg polyplexes
Amplitude (μV)
Time (sec)

N

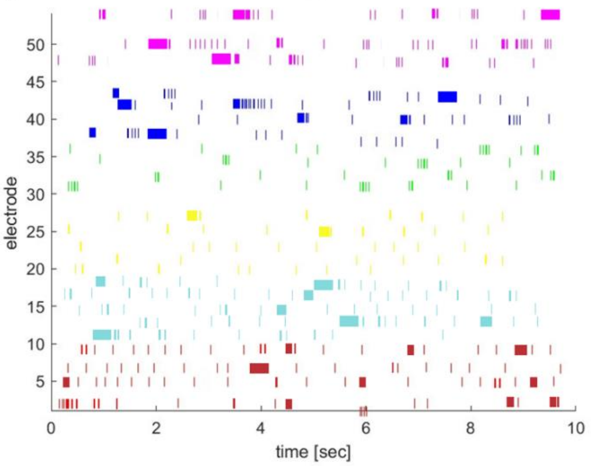


O

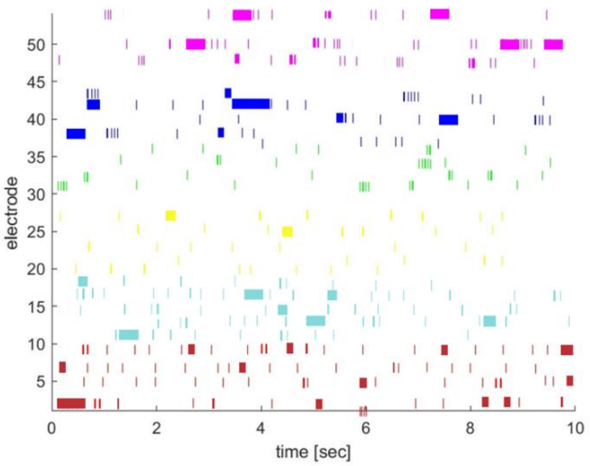


**Figure S4.** Spontaneous electrical activity of (A-F) motor and (G-L) cortical neurons cultured recorded with microelectrode arrays (MEAs) in well plates after incubation with nTg or Tg polyplexes carrying siPTEN (DIV14). Cells treated with Triton™ X-100, 0.1% (v/v) in PBS 1× were used as controls. (A) and (G) Number of active electrodes. Active electrodes were defined by showing 15 or more spikes per minute; (B) and (H) Number of spikes per minute; (C) and (I) number of bursts during 5-min recording. Bursts were defined as three or more spikes with a 300-ms interspike interval (ISI); (D) and (J) number of spikes in burst; (E) and (K) ISI per active electrode; (F) and (L) mean intra-burst interval (IBI). Results are represented as the mean of three independent experiments (n = 3). For statistical analysis, a two-way ANOVA test was used. No statistically significant differences were identified resulting from treatment with nTg or Tg polyplexes (N/P 4) compared to the non-treated (NT) cells. All graphs use a voltage-threshold based algorithm applied to 20,000 Hz high-pass filtered traces, with the threshold set to 4.5 times the standard deviation of the noise. (M) Representative spontaneous electrophysiological activity of NT neurons, neurons treated with Triton™ X-100, 0.1% (v/v) in PBS 1×, and neurons treated with Tg polyplexes (N/P 4). Raster plot of spikes from more than 20 electrodes in cultures of (N) motor neurons or (O) cortical neurons, showing the pattern and density of spikes. Results are shown as mean ± SD of three independent experiments (n = 3) with one microfluidics per experiment. For statistical analysis, two-way ANOVA test was used. Significant differences: ****$p \leq 0.0001$.

#### 5) Cellular association of non-targeted TMCSH-derives nanoparticles

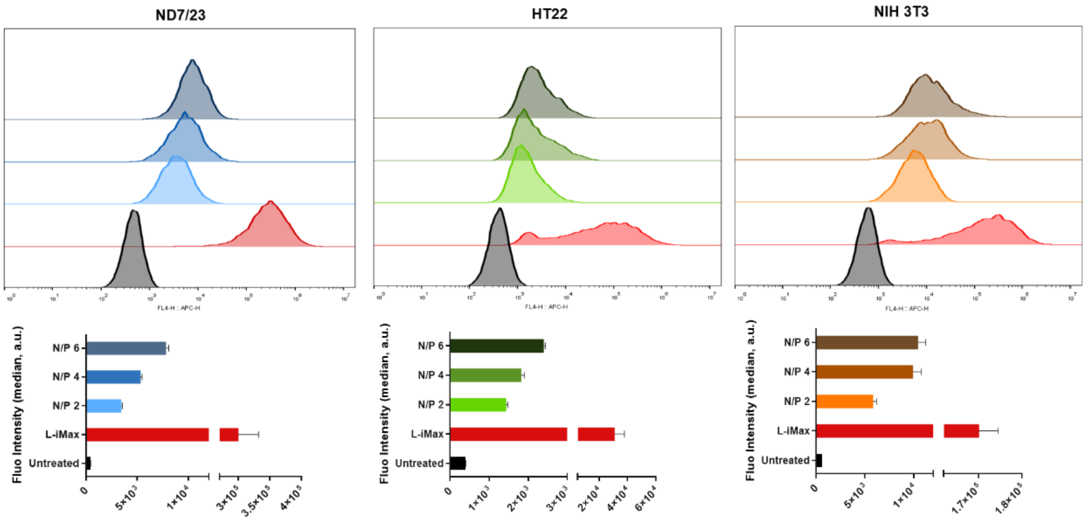


**Figure S5.** Cellular interaction of non-targeted (nTg) polyplexes. ND7/23 (in tons of blue), HT22 (in tons of green), and NIH3T3 (in tons of orange) cells were incubated for 24 hours with nTg polyplexes (N/P ratios 2-6) complexing Cy5-labeled siRNAmi (Cy5-siRNAmi) at a final siRNAmi concentration of 100 nM. Lipofectamine iMax (L-iMax) was used as a reference following the manufacturer's guidelines. Characterization and quantification were performed by flow cytometry. The coloured areas correspond to colonies of cells with high relative fluorescence (top row). Results are represented as mean ± SD of three independent experiments (n = 3)

.

#### 6) Axonal transport of neuron-targeted polyplexes – Kymographic analysis

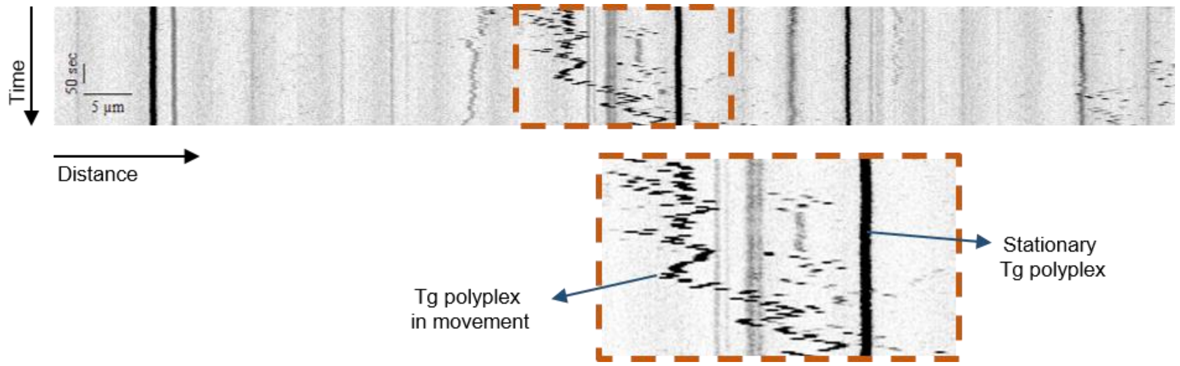


**Figure S6.** Representative kymograph showing targeted NPs pausing or moving within the axons of motor neurons.

#### 7) Biocompatibility profile of polyplexes assessed in microfluidic neuronal cultures

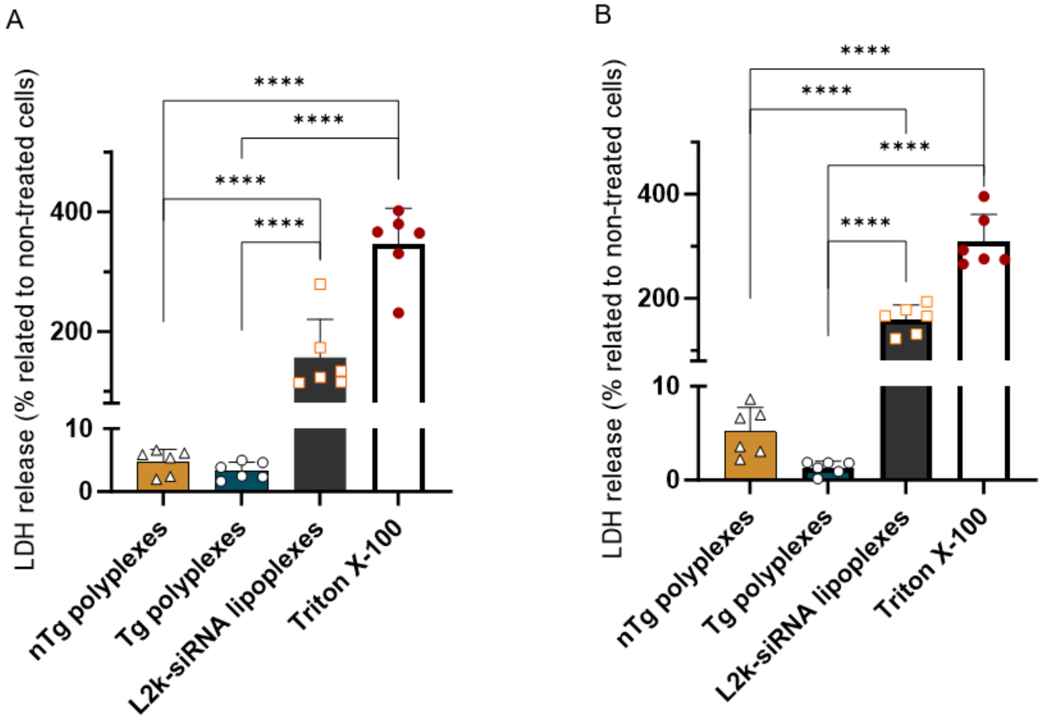


**Figure S7.** Biocompatible profile evaluation in primary motor and cortical neurons. Plasma membrane integrity of A) motor neurons and B) cortical neurons seeded in two-compartment microfluidic devices, assessed by LDH assay 24 hours incubation with N/P 4 polyplexes. Polyplexes were added in the axonal compartment and 24 hours later, culture medium from cell soma compartment was collected. Results are represented as the mean of three independent experiments (n = 3). For statistical analysis, one-way ANOVA test was used. Significant differences: ****$p \leq 0.0001$.

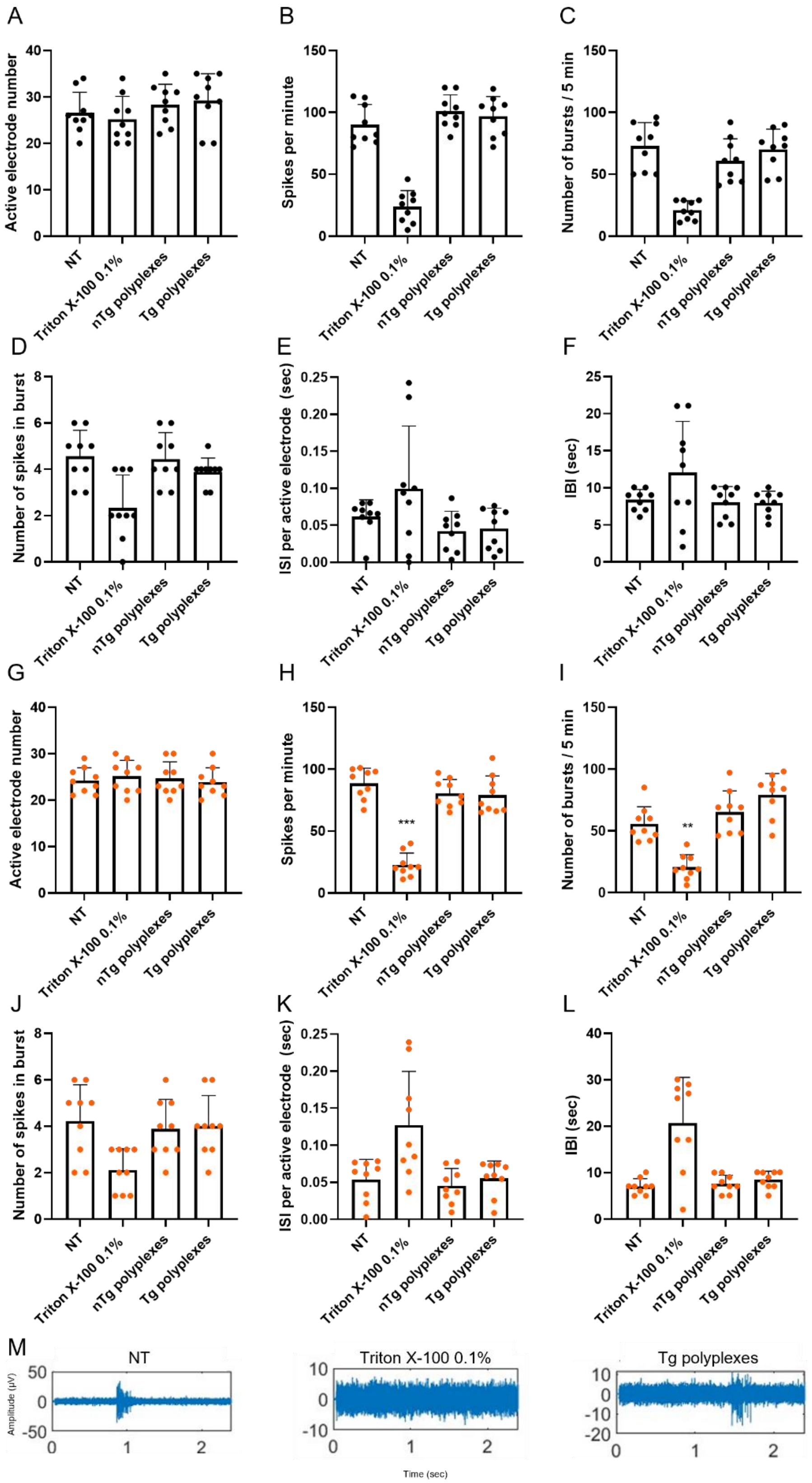


**Figure S8.** Spontaneous electrical activity of (A-F) motor and (G-L) cortical neurons, cultured in two-compartment microfluidic platforms, recorded with microelectrode arrays (MEAs) after incubation with nTg or

Tg polyplexes carrying siPTEN (DIV14). Untreated cells and cells treated with Triton™ X-100, 0.1% (v/v) in PBS 1× were used as controls. (A) and (G) Number of active electrodes. Active electrodes were defined by showing 15 or more spikes per minute; (B) and (H) Number of spikes per minute; (C) and (I) number of bursts during 5-min recording. Bursts were defined as three or more spikes with a 300-ms interspike interval (ISI); (D) and (J) number of spikes in burst; (E) and (K) ISI per active electrode; (F) and (L) mean intra-burst interval (IBI). Results are represented as the mean of three independent experiments (n = 3). For statistical analysis, a two-way ANOVA test was used. No statistically significant differences were identified resulting from treatment with nTg or Tg polyplexes (N/P 4) compared to the non-treated (NT) cells. A voltage-threshold based algorithm applied to 20,000 Hz high-pass filtered traces was used, with the threshold set to 4.5 times the standard deviation of the noise. (M) Representative spontaneous electrophysiological activity of NT neurons, neurons treated with Triton™ X-100, 0.1% (v/v) in PBS 1×, and neurons treated with Tg polyplexes (N/P 4). Results are shown as mean ± SD of three independent experiments (n = 3) with one microfluidics per experiment. For statistical analysis, two-way ANOVA test was used. Significant differences: $^{**}p < 0.01$ and $^{***} p < 0.001$.
